# L-2-hydroxyglutarate impairs neuronal differentiation through epigenetic activation of *MYC* expression

**DOI:** 10.1101/2025.04.16.649033

**Authors:** Wen Gu, Xun Wang, Ashley Solmonson, Ling Cai, Yi Xiao, Alpaslan Tasdogan, Jordan Franklin, Yuannyu Zhang, Hua Zhang, Aundrea K. Westfall, Ashley Rowe, Hetali Trivedi, Brandon Faubert, Zheng Wu, Jessica Sudderth, Lauren G. Zacharias, Bushra Afroze, Ilya Bezprozvanny, Sunil Sudarshan, Feng Cai, Samuel K. McBrayer, Thomas P. Mathews, Ralph J. DeBerardinis

## Abstract

High levels of L- and D-2-hydroxyglutarate, the reduced forms of α-ketoglutarate (αKG), are implicated in human neurodevelopmental disorders and cancer. Both enantiomers exert effects on epigenetics by modulating a family of αKG-dependent dioxygenases involved in histone, DNA and RNA demethylation. L-2HG dehydrogenase (L2HGDH) converts L-2HG to αKG. Its deficiency is a rare, autosomal recessive inborn error of metabolism (IEM) characterized by systemic elevations of L-2HG, progressive neurological disability and a high risk of malignancy in the brain. The mechanisms behind these aberrations are unknown. Here we used an isogenic, patient-derived induced pluripotent stem cell (iPSC) system to study the impact of L2HGDH deficiency on neural progenitor cell (NPC) function and neuronal differentiation. We demonstrate that L2HGDH deficiency causes accumulation of L-2HG, NPC hyperproliferation, increased clonogenicity, excessive growth, and defective neuronal differentiation in 2D cultures and cortical spheroids. Editing the *L2HGDH* locus to wild-type reverses these effects. Blocking L-2HG accumulation in NPCs with a glutaminase inhibitor also induces neuronal differentiation. L-2HG-dependent inhibition of KDM5 histone demethylases leads to widespread retention of H3K4me2 and H3K4me3, markers of active gene expression. These marks are prominently elevated at the *MYC* locus in L2HGDH-deficient cells, and consequently cells express high *MYC* both in 2D culture and in many distinct cell types within cortical spheroids. Although thousands of loci display altered histone methylation, genetically or pharmacologically normalizing *MYC* is sufficient to completely reverse defective neuronal differentiation. These data indicate that the primary metabolic disturbance in an iPSC IEM model activates the *MYC* oncogene, favoring stem cell self-renewal and suppressing lineage commitment to neurons.

## INTRODUCTION

L-2-hydroxyglutaric aciduria (L-2HGA) is a rare autosomal recessive inborn error of metabolism (IEM) caused by mutations in *L2HGDH*(1), the gene encoding L-2-hydroxyglutarate dehydrogenase, an oxidoreductase that interconverts L-2HG and α-ketoglutarate(2). Individuals with L-2HGA present with developmental delay, seizures, and progressive neurological decline(3), along with an elevated risk of tumors in the central nervous system (CNS)(4) and elsewhere(5). These neurodevelopmental abnormalities and malignancies imply altered cellular differentiation and proliferation.

Despite the identification of L-2HG as a hallmark metabolite in these patients, the underlying mechanisms connecting its accumulation to impaired neuronal function are poorly understood. Unlike many other IEMs associated with neurodevelopmental disabilities, patients with L-2HGA do not display acute encephalopathic crises or overt signs of systemic metabolic dysfunction, such as acidosis, hypoglycemia, or hyperammonemia.

2-hydroxyglutarate (2-HG) exists in two enantiomeric forms, L-2HG and D-2HG, each with distinct biological and pathological effects. Most mechanistic evidence connecting 2-HG to disease stems from studies on D-2HG in cancer, particularly in gliomas(6) and acute myeloid leukemia (AML)(7). D-2HG accumulates as a result of mutations in isocitrate dehydrogenase-1 or -2 (IDH1, IDH2)(8, 9), which confer a neomorphic enzymatic activity that converts α-ketoglutarate (αKG) to D-2HG. This accumulation inhibits αKG-dependent dioxygenases(10), including histone demethylases(11) and the Ten-eleven translocation (TET) family of hydroxylases that promote DNA demethylation(12). This enzymatic inhibition drives hypermethylation, promoting tumorigenesis and blocking differentiation. These findings have established D-2HG as an oncometabolite and a central player in epigenetic reprogramming associated with malignancy.

Elevated L-2HG levels have been observed in a variety of contexts, including in clear cell renal cell carcinoma (ccRCC) due to reduced L2HGDH activity(13, 14), as well as in metabolic stress scenarios such as hypoxia(15–17), acidosis(18), and electron transport chain (ETC) dysfunction(19). L-2HG accumulation is also observed in other IEMs, such as lipoyltransferase-1 deficiency(20). This metabolite is increasingly recognized for its effects on development and regenerative processes outside of cancer. In mouse blastocysts, elevated L-2HG levels inhibit histone demethylases, resulting in widespread histone methylation and transcriptional dysregulation after fertilization(21).

During larval growth in fruit flies, timed accumulation and degradation of L-2HG is necessary for proper epigenetic regulation of gene expression, suggesting a developmental role for this metabolite(22). The effects of L-2HG appear highly context-dependent. For example, glial-derived L-2HG promotes axon regeneration by acting on neuronal metabotropic GABAB receptors to enhance cAMP signaling. Conversely, increased L-2HG within neurons negatively regulates neuroregeneration in the *Drosophila* CNS(23).

Despite these advances, the mechanisms by which L-2HG interferes with neuronal function in humans remain poorly understood. Here we investigate the impact of L-2HG on neuronal differentiation using patient-derived, isogenic neural progenitor cells (NPCs) with either functional or deficient L2HGDH. We conduct metabolic, transcriptional, and epigenetic characterization, alongside functional perturbations and analyses in both 2D and organoid neuronal differentiation models. In so doing, we reveal the metabolic and functional consequences of L-2HG excess during neuronal specification, identify the molecular mechanism driving these abnormalities, and demonstrate improved neuronal differentiation and function upon reversal of L-2HG’s effects on key epigenetic targets.

## RESULTS

### Genomic and metabolic analysis of patients with L-2HGA

We identified a homozygous loss of function variant in *L2HGDH* (c.829C>T; p.Arg277Ter) in two siblings diagnosed with L-2HGA in Pakistan (Figure 1A, Patients 1 and 2). We confirmed that both siblings inherited one variant from each heterozygous carrier parent, consistent with the known autosomal recessive inheritance of this disease(2). Metabolomic profiling of plasma revealed elevated 2HG levels in the patients relative to their parents and control subjects (Figure 1B). Metabolite differential analysis demonstrated that 2HG was by far the most significantly altered metabolite in the plasma of the L-2HGA patients (Figure 1C). We also performed metabolomics in fibroblast cultures from both siblings and observed marked 2HG accumulation compared to fibroblasts from 30 controls (Figure 1D Supplemental Figure 1A). We then quantified the levels of both 2HG enantiomers in fibroblasts from the patients and controls; this confirmed elevated L-2HG in the patients’ cells, with no accumulation of D-2HG, as expected (Figure 1E and S1B).

**Figure 1.**
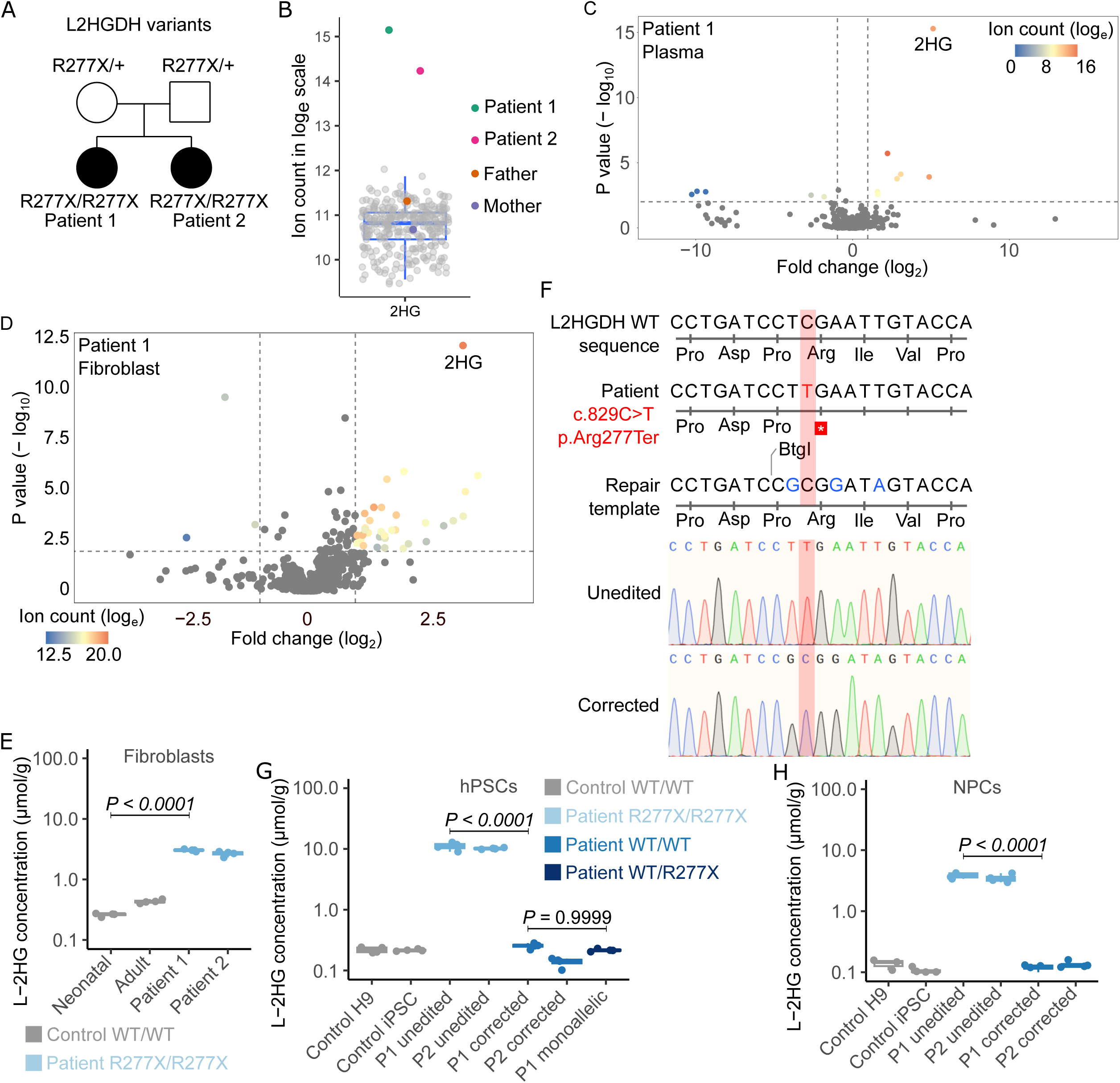
An isogenic, patient-derived iPSC system to study L2HGDH deficiency. (A) *L2HGDH* genotypes of two L-2HGA patients and their parents. (B) Relative 2HG abudnace in plasma from L-2HGA patients, parents, and unrelated subjects. (C) Volcano plot of plasma metabolites comparing Patient 1 to unrelated subjects (n = 427; sibling excluded). A linear mixed-effects model was used to account for repeated measures per individual, with subject ID as a random effect. P-values were adjusted for multiple testing (Benjamini-Hochberg), with significance thresholds of adjusted p < 0.01 and fold-change > 2 or < 0.5. (D) Volcano plot of fibroblast metabolites comparing Patient 1 to 29 unrelated lines (sibling excluded), each profiled in quadruplicate. A mixed-effects model was used as in (C), with cell line as the random effect. (E) L-2HG concentrations in neonatal control, adult control, and patient fibroblasts. (F) Alignment showing the pathogenic L2HGDH variant (c.829C>T, red), and edits (blue) introduced by the single-stranded oligonucleotide (ssODN) repair template. Electropherograms show unedited and biallelically corrected iPSC alleles. (G and H) L-2HG levels in PSCs (G) and NPCs (H) from control and patient lines with unedited or corrected L2HGDH alleles. For (B), (E), (G), and (H), data are shown as box-and-whisker plots with jittered points (n = 4 biological replicates for E–H); boxes represent the 25th–75th percentile, horizontal lines indicate medians, and whiskers extend to 1.5× the interquartile range. Statistical significance was assessed using one-way ANOVA with Tukey’s HSD test.

### CRISPR/Cas9-mediated correction of L2HGDH mutation

To model L-2HGA in culture and explore the molecular basis of neuronal dysfunction, we generated induced pluripotent stem cells (iPSCs) from patient fibroblasts using the reprogramming factors OCT4, KLF4, SOX2, and c-MYC introduced via a non-replicating Sendai virus (SeV)(24). Eight clonal cell lines were derived from the reprogrammed fibroblasts for each donor, and two of these lines from each donor were further expanded and characterized. We then applied CRISPR/Cas9-mediated genome editing to correct the c.829C>T mutation in patient-derived iPSCs, using a single-stranded oligo DNA nucleotide (ssODN) repair template (Figure 1F and Supplemental Figure1C). Sanger sequencing confirmed correction of *L2HGDH* in patient-derived iPSC clones, demonstrating precise incorporation of blocking mutations (in blue) to prevent Cas9 re-cutting(25). In subsequent experiments, “corrected” clones are ones in which *L2HGDH* was edited to the wild-type sequence, and “unedited” clones are ones that were subjected to the editing workflow but did not become edited (Supplemental Figure 1D). We measured L-2HG concentrations in pluripotent stem cells from control subjects and from corrected and unedited iPSCs from both patients (Figure 1G). Unedited iPSCs retained high L-2HG levels, but edited iPSCs had L-2HG levels comparable to control hPSCs. Consistent with autosomal recessive inheritance of L-2HGA, monoallelic homologous-directed recombination reduced intracellular L-2HG to levels comparable to biallelically corrected iPSCs (Figure 1G). We then differentiated these iPSCs into neural progenitor cells (NPCs) according to a well-established dual SMAD inhibition neural specification protocol (26) and found that unedited NPCs from both patients had high L-2HG levels but corrected NPCs had low L-2HG levels comparable to NPCs from controls (Figure 1H). D-2HG levels were comparable in all these cell lines (Supplemental Figure 1E).

### *L2HGDH*-mutant NPCs are hyperproliferative with enhanced self-renewal capacity and impaired differentiation to neurons

Unedited NPCs were highly proliferative, exceeding the proliferation rates of corrected NPCs and control H9 NPCs (Figure 2A). We generated cortical spheroids from H9, unedited and corrected NPCs, because these 3-dimensional models of differentiation can be used to study disease processes that impact the brain(27). After 30 days of differentiation, spheroids derived from unedited NPCs were much larger than those from corrected and control NPCs (Figure 2B). Quantification of spheroid surface area confirmed the increased size of unedited relative to corrected spheroids, whereas no significant difference was detected between corrected and control spheroids (Figure 2C). Next, to evaluate NPC self-renewal capacity, we cultured single NPCs and measured their ability to form colonies(28). Stereoscopic and phase contrast imaging revealed enhanced colony formation in unedited NPCs compared to corrected cells (Figure 2D). Quantification of sphere-forming frequency demonstrated a large increase in the proportion of sphere-forming cells in unedited NPCs relative to corrected lines (Figure 2E).

**Figure 2.**
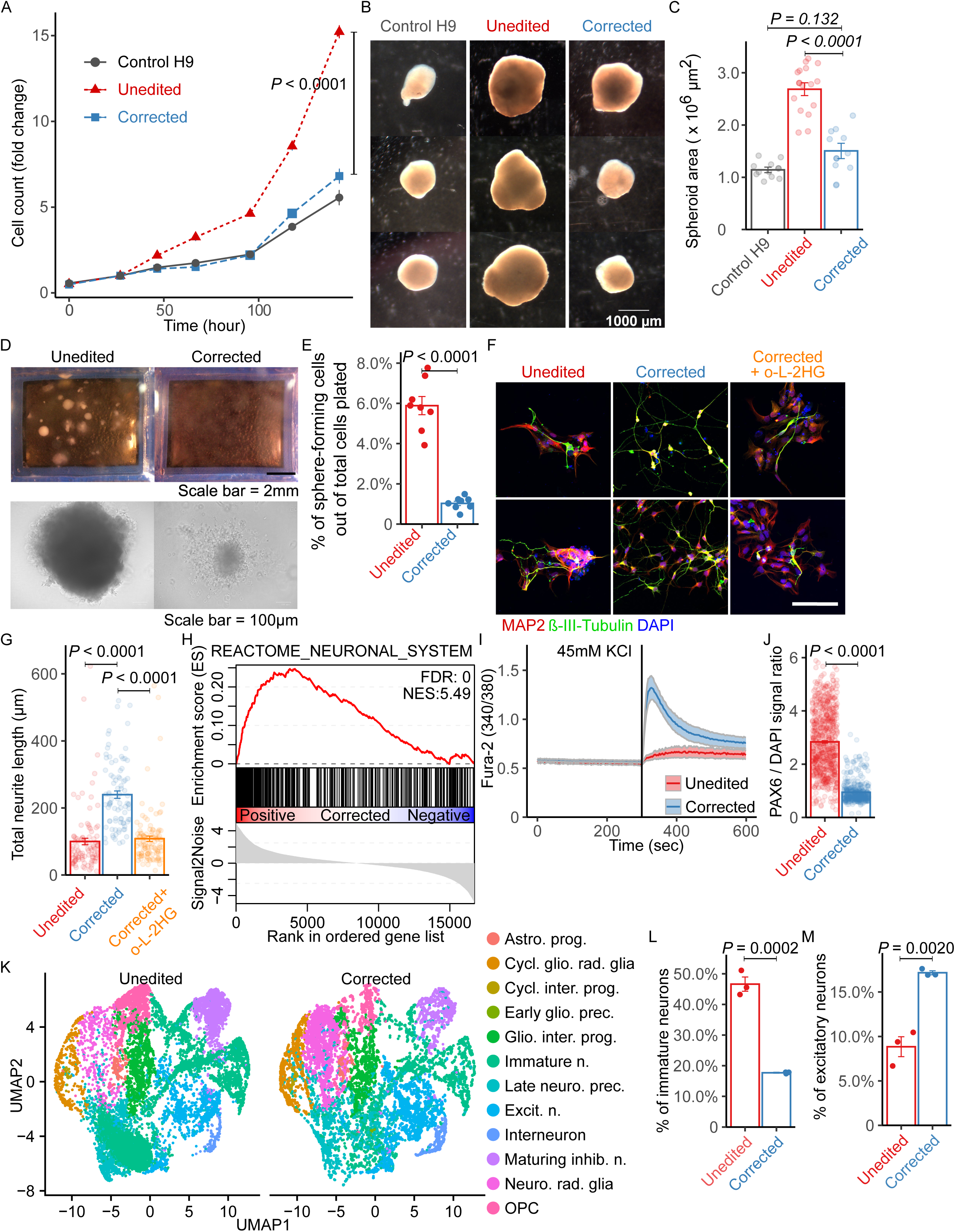
L-2HG accumulation enhances NPC proliferation and self-renewal while inhibiting neuronal differentiation. (A) Growth curves of NPCs derived from H9 control, Patient 1 unedited, and Patient 1 corrected iPSCs. (B) Stereoscopic images of cortical spheroids at day 30 of differentiation. Scale bar, 1000 µm. (C) Quantification of spheroid surface area at day 30. (D) Images of colonies formed from single NPCs (top: stereoscope; bottom: phase contrast). Scale bars, 2 mm and 100 µm. (E) Frequency of colony formation in single NPCs. (F) Immunostaining for MAP2 and βIII-Tubulin in Patient 1 unedited, corrected, and corrected + 30 µM octyl-L-2HG (o-L-2HG) neurons at day 14. DAPI marks nuclei. Scale bar, 100 µm. (G) Quantification of neurite length from images in (F) using the SNT plugin in ImageJ. (H) GSEA plot showing enrichment of the Reactome_Neuronal_System gene set in corrected versus unedited neurons. (I) Fura-2 ratio recordings of Ca²⁺ dynamics during 45 mM KCl application in neurons differentiated for 47 days (unedited: n = 29; corrected: n = 24). (J) Quantification of PAX6 signal intensity normalized to DAPI in unedited and corrected neurons. (K) UMAP of single-cell transcriptomes from day 45 cortical spheroids (unedited and corrected; 12,660 cells each). Cell type annotations include: astrocyte progenitor (Astro. prog.), cycling gliogenic radial glia (Cycl. glio. rad. glia), cycling intermediate progenitor (Cycl. inter. prog.), early gliogenic precursor (Early glio. prec.), gliogenic intermediate progenitor (Glio. inter. prog.), immature neuron (Immature n.), late neurogenic precursor (Late neuro. prec.), excitatory neuron (Excit. n.), interneuron, maturing inhibitory neuron (Maturing inhib. n.), neurogenic radial glia (Neuro. rad. glia), and oligodendrocyte progenitor cell (OPC). (L and M) Proportions of immature neurons (L) and excitatory neurons (M) among cell types shown in (K). For (A), (C), (E), (G), (J), (L), and (M), data are presented as mean ± 1 SEM. For (A), differences in cell growth were analyzed using two-way ANOVA (cell line × time), followed by Tukey’s HSD post hoc test. Statistical significance was assessed using Student’s t-test for (E) (n = 8), (J) (n = 679 unedited; n = 430 corrected), and for (L) and (M) (n = 3 each). One-way ANOVA followed by Tukey’s HSD test was used for (C) (n = 10 H9, n = 15 unedited, n = 10 corrected) and (G) (n = 96 unedited, n = 86 corrected, n = 108 corrected + o-L-2HG).

Next, we investigated the effects of *L2HGDH* mutation on neuronal differentiation by subjecting patient-derived NPCs to a well-described neuronal differentiation protocol(29). This approach generates a majority of VGLUT1-positive neurons, which are presumed to be glutamatergic forebrain neurons(26). We immune-stained day 14 two-dimensional neuronal cultures with MAP2 and β-III-Tubulin, markers of mature neurons. Morphologically, the corrected cells displayed extensive neurite outgrowth, characteristic of neurons generated in this assay, whereas neurite formation was blunted in the unedited cells (Figure 2, F and G). Supplementing corrected NPCs with a membrane-permeable L-2HG ester(10) (30 µM octyl-L-2HG) completely reversed the effect of *L2HGDH* correction on total neurite length (Figure 2, F and G). This indicates an L-2HG-dependent defect in neurite growth in the unedited cells.

RNA-sequencing was performed to further characterize the molecular differences between unedited and corrected cells. Gene set enrichment analysis (GSEA) indicated enrichment of the REACTOME_NEURONAL_SYSTEM gene set in corrected compared to unedited neurons after 14 days of differentiation (Figure 2H); this was the highest-scoring of all 3917 gene sets by GSEA (Supplemental Figure 2A and Supplemental Table 1). Furthermore, to assess neuronal excitability, functional calcium imaging was performed in neuronal cultures following KCl stimulation(30). This assay revealed essentially no response in most unedited NPC-derived neurons, but prominent intracellular calcium increases in the corrected cells, indicative of improved neuronal activity (Figure 2I). Consistent with a delayed exit from the progenitor state, unedited neurons retained much higher levels of PAX6, a well-established marker of neural progenitor cells, compared to their corrected counterparts after 14 days of neuronal differentiation (Figure 2J).

To further resolve cell-type composition during neuronal differentiation in 3D, we performed single-cell RNA sequencing on day 45 cortical spheroids derived from unedited and corrected hiPSCs. UMAP embedding revealed distinct differences in cellular composition between conditions (Figure 2K). Unedited spheroids exhibited a increased proportion of immature, uncommitted neurons and a corresponding reduction in specified excitatory neurons relative to corrected spheroids (Figure 2, L and M), consistent with a block in neuronal specification.

Together, these findings highlight the impact of *L2HGDH* mutation and L-2HG accumulation on NPC proliferation, self-renewal, and neuronal differentiation. Our data demonstrate that CRISPR/Cas9-mediated correction of L2HGDH restores neuronal differentiation and function, suggesting the potential of this isogenic model to uncover the molecular mechanism by which L-2HG excess disrupts neuronal development.

### Depleting L-2HG enhances neuronal differentiation in L2HGDH-deficient NPCs

We next sought to test whether the defect in neuronal differentiation in L2HGDH-deficient NPCs could be reversed pharmacologically. We identified the predominant nutrient source of L-2HG by performing parallel stable isotope tracing experiments in medium containing [U-^13^C]glucose and unlabeled glutamine, or [U-^13^C]glutamine and unlabeled glucose. [U-^13^C]glucose is predicted to give rise to αKG and L-2HG m+2, while [U-^13^C]glutamine is predicted to give rise to αKG and L-2HG m+5 (Figure 3A; complete set of labeling data in Supplemental Table 2 and Supplemental Table 3).

**Figure 3.**
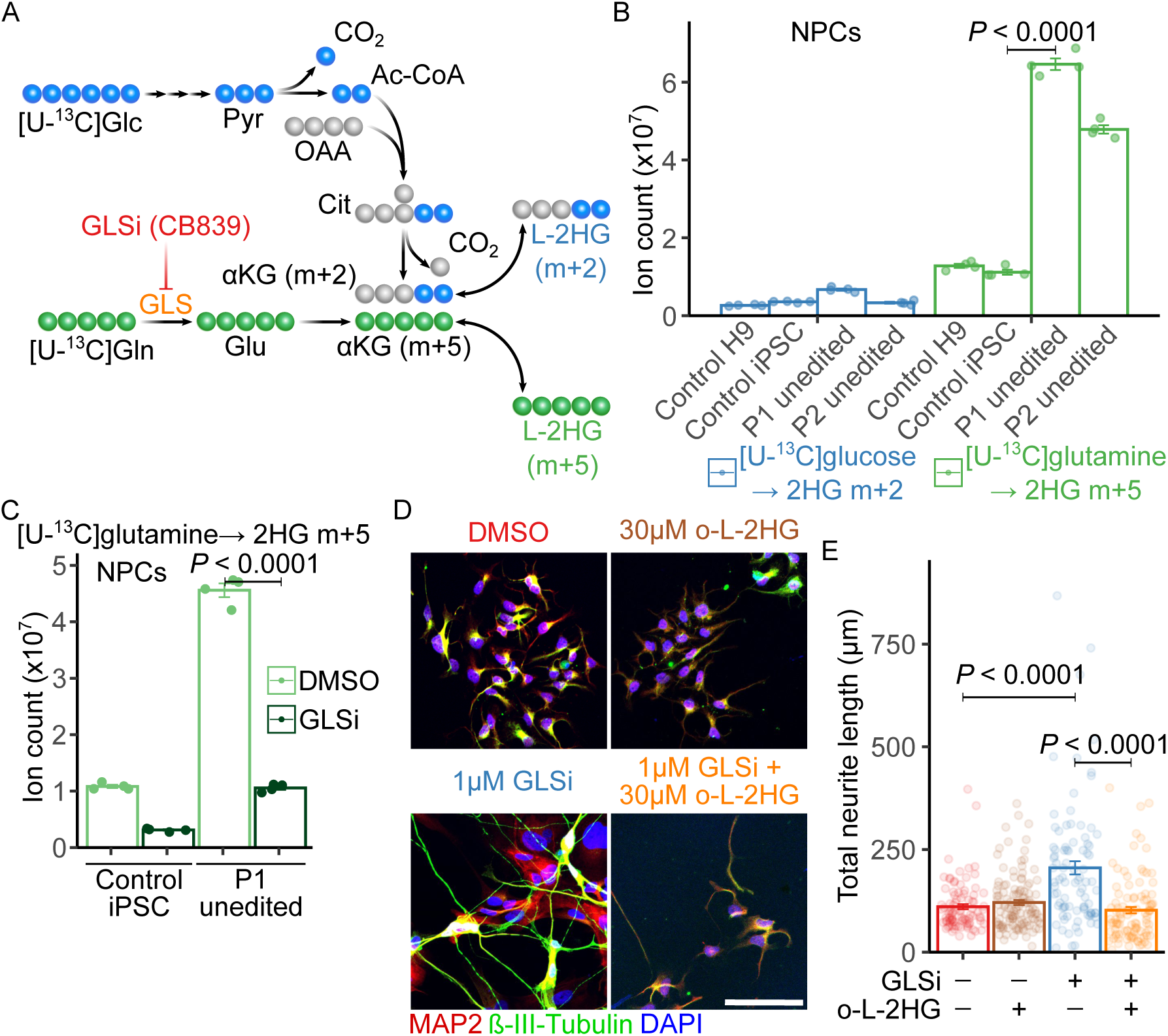
Suppressing L-2HG synthesis improves neurite formation in L2HGDH-deficient NPCs. (A) Schematic of ¹³C-labeling routes to L-2HG from [U-¹³C]glucose (blue) or [U-¹³C]glutamine (green) in NPCs. Abbreviations: Glc, glucose; Pyr, pyruvate; Ac-CoA, acetyl-CoA; OAA, oxaloacetate; Cit, citrate; αKG, α-ketoglutarate; Gln, glutamine; GLS, glutaminase; GLSi, glutaminase inhibitor (CB839); Glu, glutamate. (B) Ion counts of ¹³C-labeled 2HG isotopologues in NPCs derived from H9, control hiPSC, and Patients 1 and 2 after 4 h of labeling with [U-¹³C]glucose or [U-¹³C]glutamine. (C) Ion counts of ¹³C-labeled 2HG isotopologues in control hiPSC-derived and Patient 1 NPCs cultured in [U-¹³C]glutamine and treated with DMSO or 1 µM CB839 (GLSi) for 4 h. (D) Immunofluorescence for MAP2 and βIII-Tubulin in Patient 1 NPCs after 14 days of neuronal differentiation, treated with DMSO, 30 µM octyl-L-2HG (o-L-2HG), 1 µM CB839 (GLSi), or both. DAPI marks nuclei. Scale bar, 100 µm. (E) Quantification of total neurite length from cells in (D), measured with the SNT plugin in ImageJ. For (B) and (C), ion counts were normalized to total ion current (TIC). Data were non-normally distributed (Shapiro–Wilk test), so statistical significance was assessed by Kruskal–Wallis test with Dunn’s post hoc correction (n = 4 per group). For (E), one-way ANOVA with Tukey’s HSD test was used (n = 90 for DMSO + DMSO; n = 116 for DMSO + o-L-2HG; n = 89 for GLSi + DMSO; n = 101 for GLSi + o-L-2HG). Data shown as mean ± 1 SEM.

These isotope tracing experiments used a mass spectrometry approach that does not discriminate between L- and D-2HG, but note that L-2HG accounts for most of the total 2-HG pool in L2HGDH-deficient cells (Figure 1H and Supplemental Figure 1E). In all cell lines examined, the abundance of ^13^C-labeled 2HG from glutamine exceeded the abundance from glucose, as expected if glutamine is the dominant source of αKG (Figure 3B). Unedited patient-derived NPCs cultured with [U-^13^C]glutamine had much more ^13^C-2HG than these same NPCs cultured in [U-^13^C]glucose, or control NPCs cultured in [U-^13^C]glutamine (Figure 3B).

We next investigated whether blocking glutamine metabolism could reduce L- 2HG accumulation by treating unedited NPCs with the glutaminase inhibitor CB839 (GLSi) or DMSO. GLS inhibition reduced ^13^C-labeled L-2HG (m+5) derived from [U-^13^C]glutamine, indicating that GLS is required for glutamine-derived 2HG production in both control and L2HGDH-deficient NPCs in culture (Figure 3C).

Next, we evaluated whether CB839 could enhance neuronal differentiation in L2HGDH-deficient NPCs. Immunofluorescence staining for the neuronal markers MAP2 and β-III-Tubulin at day 14 of differentiation revealed approximately a doubling of total neurite length in unedited NPCs from Patient 1 (Figure 3, D and E). This effect was completely reversed by supplementing the medium with 30 µM octyl-L-2HG (Figure 3, D and E). Octyl-L-2HG had no suppressive effect on neurite length in unedited cells treated with DMSO, where the intracellular L-2HG concentration is already high (Figure 3E). These results provide proof of concept that targeting cellular metabolism to reduce L-2HG levels can restore neuronal differentiation in L2HGDH-deficient NPCs.

### Increased *MYC* expression in L2HGDH-deficient NPCs and cortical spheroids

To broadly assess the molecular consequences of L2HGDH deficiency in NPCs and better understand the defect in neuronal differentiation, we conducted bulk RNA-seq in control, unedited and corrected NPCs. Unbiased gene set enrichment analysis (GSEA)(31) identified the upregulation of MYC target genes in unedited NPCs (NES = 5.30) (Figure 4A). Among the 3,494 gene sets examined, two MYC target gene sets ranked within the top 10 and four ranked within the top 30 in unedited NPCs (Supplemental Figure 3A and Supplemental Table 4). RNA sequencing also revealed a marked increase in *MYC* mRNA levels in unedited patient NPCs compared to both corrected NPCs and control H9 NPCs (Figure 4B). Immunoblot analysis of nuclear lysates revealed elevated c-MYC in unedited NPCs relative to corrected NPCs, supporting the transcriptional data (Figure 4C).

**Figure 4.**
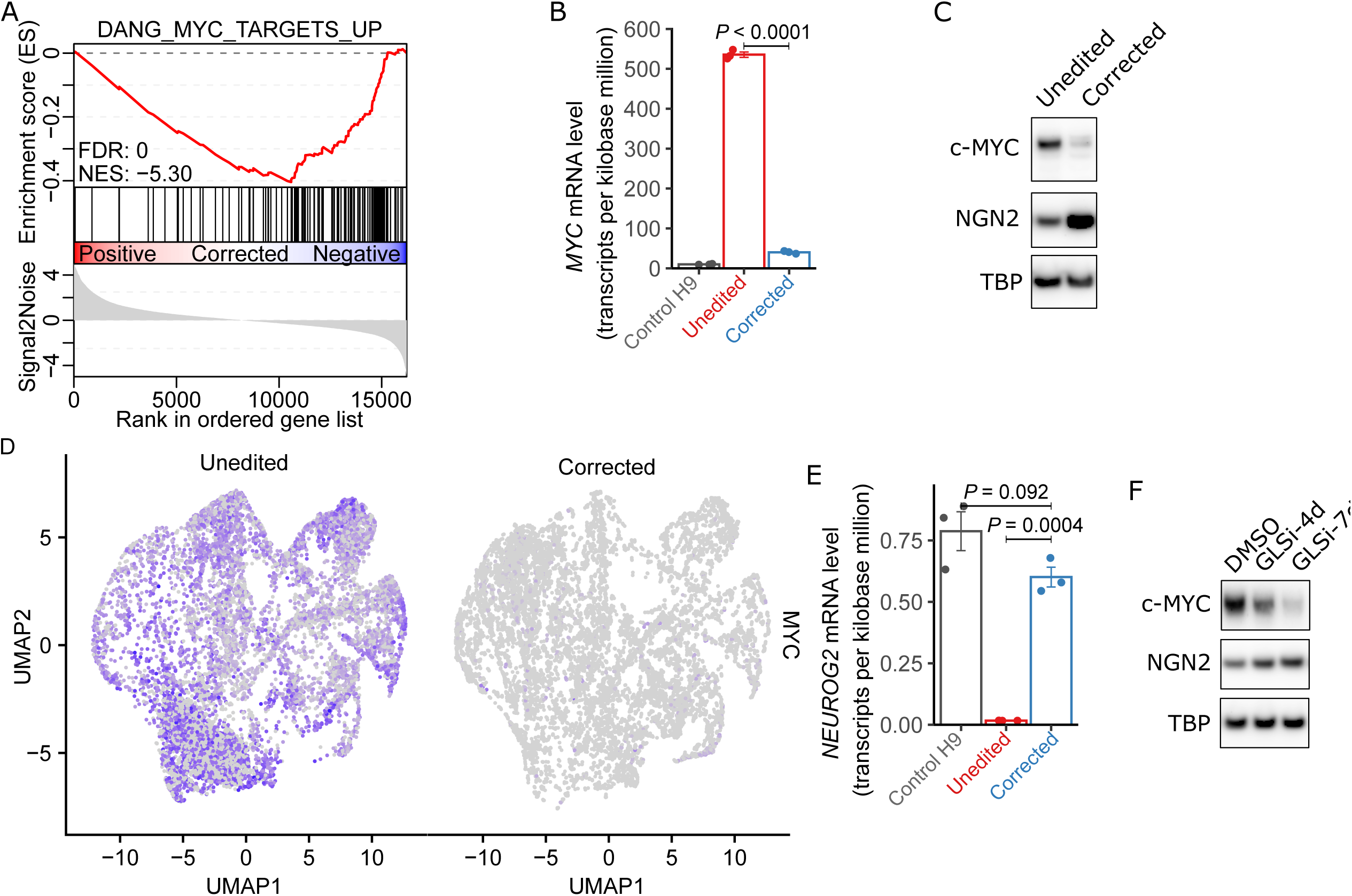
Increased *MYC* expression in L2HGDH-deficient NPCs and cortical spheroids. (A) Gene set enrichment analysis (GSEA) mountain plot showing enrichment of Dang_MYC_Targets_Up gene set in unedited versus corrected Patient 1 NPCs. (B) Quantification of *MYC* mRNA in control H9 NPCs, unedited Patient 1 NPCs, and corrected Patient 1 NPCs. mRNA levels were obtained from RNA sequencing of biological triplicates and expressed as transcripts per kilobase million (TPM). (C) Immunoblot analysis of nuclear c-MYC and Neurogenin-2 in unedited and corrected Patient 1 NPCs. TBP was used as a loading control for nuclear lysates. (D) UMAP heatmaps displaying MYC expression in single-cell RNA-seq of unedited and corrected day 45 cortical spheroids. Color intensity (purple) indicates *MYC* transcript levels in individual cells. (E) Quantification of *NEUROG2* mRNA in control H9 NPCs, unedited Patient 1 NPCs, and corrected Patient 1 NPCs. mRNA levels were obtained from RNA sequencing of biological triplicates and expressed as transcripts per kilobase million (TPM). (F) Immunoblot analysis of nuclear c-MYC and Neurogenin-2 in Patient 1 NPCs treated with DMSO or 1 µM CB839 (GLSi) for 4 or 7 days. TBP served as a loading control for nuclear lysates. For (B) and (E), statistical significance was assessed on log-transformed TPM values using one-way ANOVA followed by Tukey’s HSD test. Error bars represent ±1 SEM of three biological replicates.

We also evaluated MYC expression in day 45 cortical sphperoids and observed a marked increase in *MYC* transcript levels in unedited spheroids relative to corrected spheroids, visualized by UMAP-based expression heatmaps (Figure 4D). *MYC* was upregulated in essentially all cell types in unedited spheroids, indicating widespread overexpression (Supplemental Figure 3B). Elevated *MYC* was observed in neurogenic radial glia, with marked upregulation in cycling intermediate progenitors and cycling gliogenic radial glia. This elevated expression persisted through both neuronal and glial lineages, including late neurogenic precursors, immature neurons, and gliogenic progenitors, suggesting that L-2HG–associated dysregulation of *MYC* occurs early in lineage commitment and is sustained across multiple descendant cell types.These data confirm that MYC overexpression extends beyond 2D NPC cultures to more complex 3D models of neurodevelopment.

Interestingly, we observed a concurrent decrease in Neurogenin-2, a key regulator of neuronal differentiation(32), in unedited cells (Figure 4E). To determine if pharmacological suppression of L-2HG production could mitigate *MYC* overexpression and restore *NEUROG2* (the gene encoding Neurogenin-2) expression, we treated unedited NPCs with CB839 (GLSi) for 4 or 7 days. The drug reduced nuclear c-MYC levels in a time-dependent manner, accompanied by an increase in Neurogenin-2 expression (Figure 4F).

To validate the increased c-MYC levels in NPCs within the embryonic mouse brain, we used a previously described *L2hgdh* knockout mouse model(33). We dissected the dorsal telencephalon of wild-type and *L2hgdh^-/-^* E15.5 mouse embryos and cultured neurospheres *ex vivo* to obtain NPCs. To test whether L2HGDH deficiency impacts neuronal differentiation in this system, wild-type and *L2hgdh^-/-^* NPCs were differentiated into neurons for three weeks, fixed, and immunostained for MAP2 and β-III-Tubulin. Quantification of neurite outgrowth revealed that *L2hgdh^-/-^* neurons exhibited shorter neurites than their wild-type counterparts (Supplemental Figure 3C). We then performed immunoblot analysis of nuclear c-MYC in wild-type and *L2hgdh^-/-^* NPCs.

Consistent with human NPC data, *L2hgdh^-/-^* NPCs exhibited a modest increase in c-MYC protein levels compared to wild-type NPCs (Supplemental Figure 3D and 3E).

### L2HGDH-deficient NPCs display enhanced activating histone methylation marks in *MYC* regulatory regions

Given L-and D-2HG’s potentially broad roles in blocking epigenetic demethylases that use αKG as a cofactor, we compared several aspects of the epigenetic landscape between unedited and corrected NPCs. We did not observe an increase in global DNA methylation at cytosine residues (5mC, Figure 5A) or in global N6-methyladenosine in mRNA (m6A, Figure 5B) in NPCs with L-2HG accumulation. Rather, m6A methylation was somewhat higher in corrected NPCs compared to unedited NPCs. Therefore at the genome-wide level, we do not observe global suppression of 5mC and m6A demethylation in these L2HGDH-deficient cells, despite 2HG’s ability to block TET DNA demethylases and the RNA demethylases ALKBH5 and FTO in other contexts(34, 35). We estimated the total (i.e. non-compartmentalized) intracellular concentration of L-2HG in L2HGDH-deficient NPCs to be 1,425 ± 75 µM (mean ± SEM). This concentration is several-fold higher than the reported IC₅₀ values for most histone lysine demethylases (KDMs)(36, 37). It is somewhat below the reported IC₅₀ for TET2(38), which may explain why L-2HG accumulation apparently does not impact global DNA methylation in L2HGDH-deficient NPCs.

**Figure 5.**
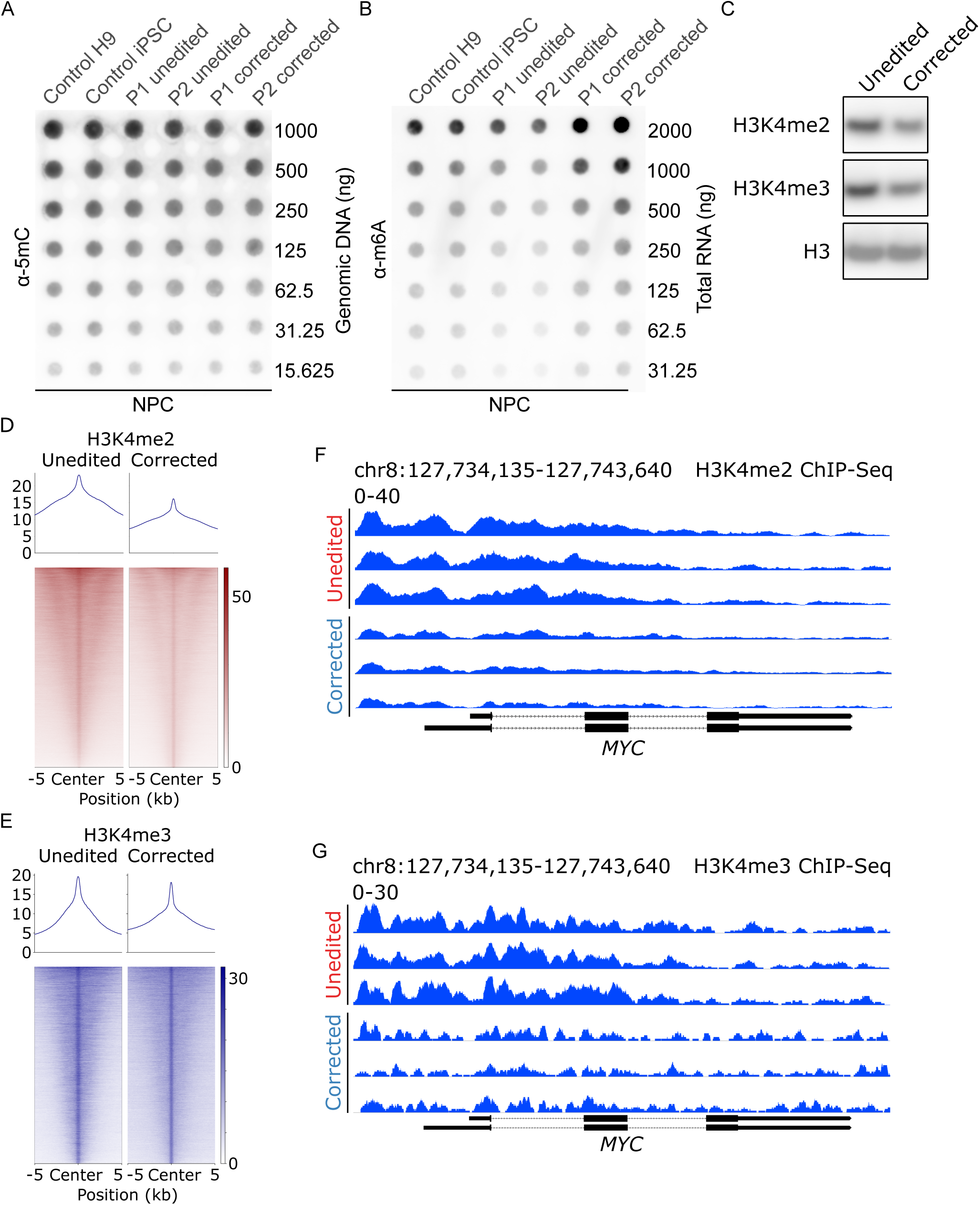
Increased activating histone methylation marks H3K4me2 and H3K4me3 at the *MYC* locus in L2HGDH-Deficient NPCs. (A) DNA dot blot analysis of 5mC levels in NPCs, including control H9 NPCs, control iPSC-derived NPCs, unedited Patient 1 NPCs, unedited Patient 2 NPCs, corrected Patient 1 NPCs, and corrected Patient 2 NPCs. (B) m6A dot blot analysis of total RNA in NPCs, including control H9 NPCs, control NPCs, unedited Patient 1 NPCs, unedited Patient 2 NPCs, corrected Patient 1 NPCs, and corrected Patient 2 NPCs. (C) Immunoblot analysis of activating histone markers H3K4me2 and H3K4me3 levels in unedited and corrected Patient 1 NPCs. Histone H3 was used as a loading control for histone lysates. (D) and (E) ChIP–Seq profiles of H3K4me2 (D) and H3K4me3 (E) in unedited and corrected Patient 1 NPCs. ChIP–Seq signals were plotted over center peaks (± 5 kb from peak center) identified in corrected Patient 1 NPCs. Sites were sorted by the ChIP–Seq signal intensity from corrected Patient 1 NPCs. (F) and (G) Representative ChIP-seq tracks showing H3K4me2 (F) and H3K4me3 (G) enrichment at the *MYC* locus in unedited and corrected Patient 1 NPCs.

We next examined histone methylation, focusing on marks associated with active transcription because of the observation that *MYC* expression is enhanced in L2HGDH-deficient NPCs. Methylation of histone H3, lysine residue 4 (H3K4) is associated with gene activation in development(39). Immunoblot analysis revealed reduced dimethylated and trimethylated H3K4 (H3K4me2 and H3K4me3) in corrected NPCs compared to unedited cells (Figure 5C). To characterize the genomic distribution of these marks, we performed chromatin immunoprecipitation sequencing (ChIP-seq) for H3K4me2 and H3K4me3. Aggregated ChIP-seq signals plotted over peak centers (±5 kb) demonstrated globally increased enrichment of both H3K4me2 and H3K4me3 in unedited NPCs relative to corrected NPCs (Figure 5, D and E). This widespread elevation suggests that L2HGDH deficiency contributes to broad epigenetic reprogramming in NPCs via changes in H3K4 methylation. Unbiased differential binding analysis of H3K4me2 and H3K4me3 ChIP-seq data further confirmed that both marks were prominently increased at *MYC* regulatory regions and ranked among the most hypermethylated sites in unedited NPCs compared to corrected NPCs (Supplemental Figure 4A, Supplemental Tables 5 and 6). Focusing on the *MYC* locus, representative ChIP-seq tracks highlighted pronounced enrichment of H3K4me2 and H3K4me3 in unedited NPCs compared to corrected cells (Figure 5, F and G). When all enriched genomic regions were mapped to their associated genes, unbiased Kyoto Encyclopedia of Genes and Genomes (KEGG) over-representation analysis (ORA) identified “Cell cycle” as the top enriched pathway for both activating H3K4 methylation marks in unedited NPCs (Supplemental Figure 4, B and C, and Supplemental Tables 7 and 8), aligning with the hyperproliferative phenotype observed in these cells. These data indicate that increased activating histone methylation at the *MYC* promoter may contribute to *MYC* overexpression in L2HGDH-deficient NPCs.

### KDM5 inhibition recapitulates the effects of L-2HG on H3K4 methylation and neuronal differentiation

KDM5 family enzymes are the primary demethylases responsible for removing H3K4 methylation(40) and are potently inhibited by L-2HG(10). Given that L2HGDH-deficient NPCs exhibit increased H3K4me2 and H3K4me3 levels, we sought to determine whether pharmacological inhibition of KDM5 would mimic the effects of L-2HG accumulation and impair neuronal differentiation. To test this, we treated corrected Patient 1 NPCs with the selective KDM5 inhibitor C70(41) (25 µM) for 14 days.

Immunoblot analysis confirmed that KDM5 inhibition increased H3K4me2 and H3K4me3 levels compared to DMSO-treated controls, supporting KDM5’s role in H3K4 demethylation in these cells (Supplemental Figure 4D). ChIP-seq analysis further revealed that KDM5 inhibition resulted in a genome-wide increase in H3K4me2 and H3K4me3 levels (Supplemental Figure 4, E and F). Representative ChIP-seq tracks confirmed that C70-treated NPCs exhibited higher H3K4me2 and H3K4me3 deposition at the *MYC* promoter compared to DMSO-treated controls (Supplemental Figure 4, G and H).

Given that L-2HG-mediated MYC overexpression is associated with impaired neuronal differentiation, we next examined whether KDM5 inhibition similarly influenced MYC levels. qPCR analysis revealed a moderate increase in *MYC* mRNA abundance in C70-treated NPCs (Supplemental Figure 4I). Similarly, immunoblot analysis of nuclear lysates revealed that C70 treatment resulted in modestly increased c-MYC expression relative to DMSO-treated NPCs (Supplemental Figure 4J). Finally, to assess the functional impact of KDM5 inhibition on neuronal differentiation, we quantified neurite outgrowth following treatment. Consistent with the effects of L-2HG accumulation, C70-treated NPCs exhibited reduced neurite length compared to controls, indicating that KDM5 inhibition negatively impacts neuronal differentiation (Supplemental Figure 4K).

These findings indicate that inhibition of KDM5, like L-2HG accumulation, increases H3K4 methylation, increases *MYC* expression, and impairs neurite length. This supports the idea that L-2HG may exert its effects, at least in part, through inhibition of KDM5 demethylases.

### c-MYC depletion restores neuronal differentiation in L2HGDH-deficient NPCs

Although correcting mutant *L2HGDH* alleles reduced H3K4 methylation at hundreds of genomic loci, the prominent effect on the *MYC* locus and the broad roles of *MYC* in cell proliferation and differentiation suggested that *MYC* expression could be sufficient to explain the effects of L2HGDH deficiency. We first ruled out the possibility that impaired neurite length was a non-specific response to enhanced proliferation.

Unedited NPCs were treated with the anti-mitotic agent nocodazole at a concentration that maintained NPC viability throughout the treatment duration (Supplemental Figure 5A). As expected, nocodazole suppressed NPC proliferation (Supplemental Figure 5B), but had no impact on the expression of c-MYC or Neurogenin-2 by immunoblot (Figure 6A). Similarly, neurite length was not enhanced by nocodazole (Figure 6B). This indicates that cell cycle modulation is insufficient to restore neuronal differentiation in L2HGDH-deficient NPCs.

**Figure 6.**
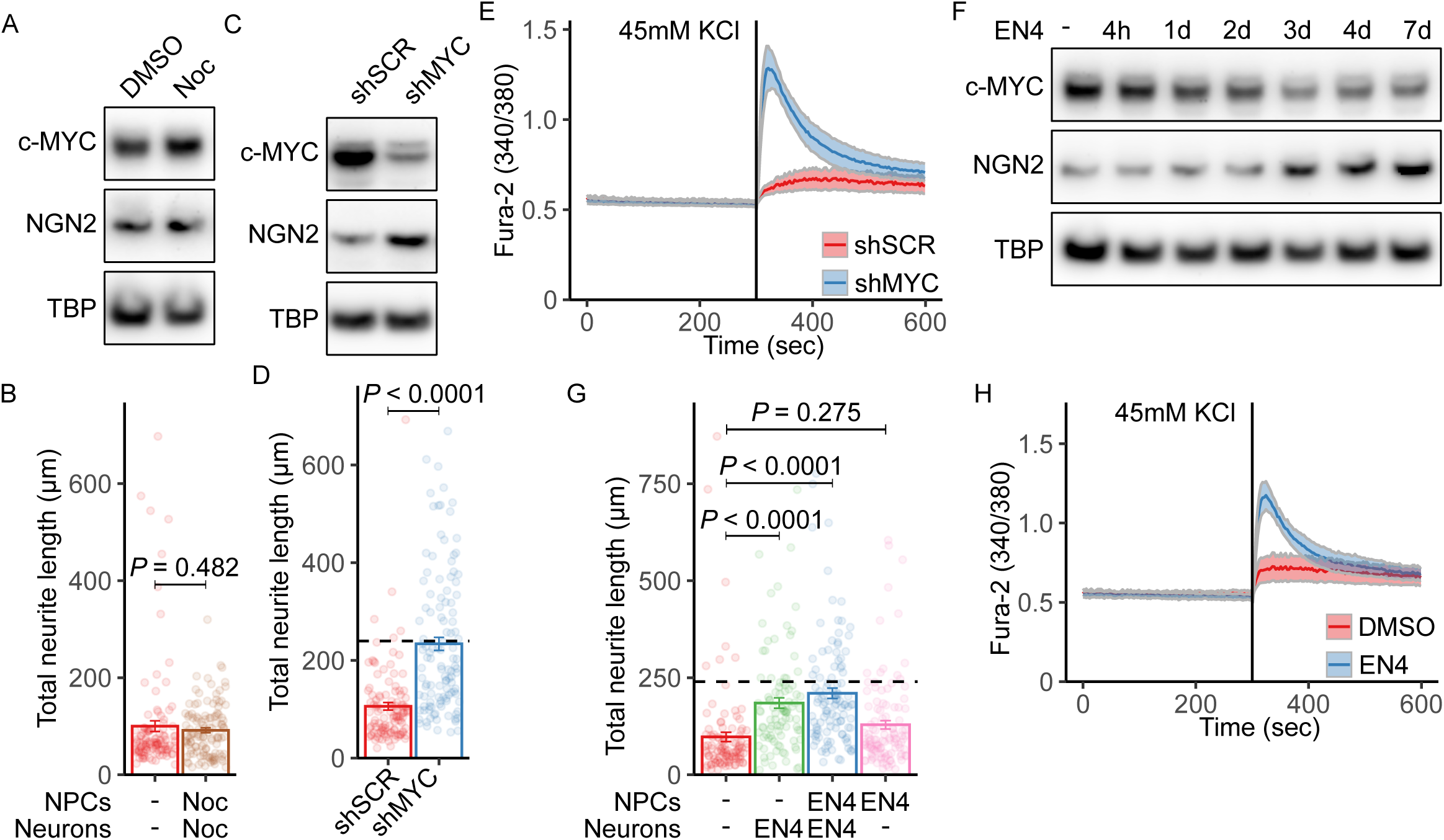
c-MYC depletion restores neuronal differentiation in L2HGDH-deficient NPCs. (A) Immunoblot of nuclear c-MYC and Neurogenin-2 in unedited Patient 1 NPCs treated with DMSO or 20 µM nocodazole (Noc) for 7 days. TBP was used as a loading control. (B) Quantification of neurite lengths in Patient 1 NPCs treated with DMSO or 20 µM Noc at the NPC stage and throughout 14-day neuronal differentiation. (C) Immunoblot of nuclear c-MYC and Neurogenin-2 in Patient 1 NPCs transduced with lentivirus expressing control shRNA (shSCR) or MYC-targeting shRNA (shMYC). (D) Quantification of neurite lengths in Patient 1 NPCs transduced with shSCR or shMYC. The black dashed line denotes the mean neurite length in corrected neurons. (E) Representative recordings of intracellular Ca²⁺ dynamics in neurons transduced with shSCR (n = 29) or shMYC (n = 24), captured every 10 s using Fura-2 ratio imaging during 45 mM KCl application. Neurons were differentiated for 47 days. (F) Immunoblot of nuclear c-MYC and Neurogenin-2 in Patient 1 NPCs treated with DMSO or 50 µM EN4 for the indicated durations. (G) Quantification of neurite lengths in Patient 1 NPCs treated with: DMSO; EN4 throughout differentiation; EN4 at the NPC stage and throughout differentiation; or EN4 only at the NPC stage. The dashed line indicates the corrected neuron mean. (H) Representative Ca²⁺ recordings in neurons treated with DMSO (n = 28) or 50 µM EN4 (n = 26), acquired as in (E). For (B), (D), and (G), neurite lengths were measured using the SNT plugin in ImageJ. Data are mean ± 1 SEM. Statistical significance was determined by Student’s t test (B, D) or one-way ANOVA with Tukey’s HSD test (G).

Next, we used lentiviral shRNA to suppress *MYC* expression. Immunoblot analysis confirmed a reduction in c-MYC protein levels in NPCs transduced with *MYC*-targeting shRNA (shMYC) lentivirus compared to a scrambled control (shSCR). This reduction was accompanied by an increase in Neurogenin-2 (Figure 6C). Quantification of neurite outgrowth showed that shMYC doubled the average total neurite length in unedited NPCs compared to shSCR controls (Figure 6D). To evaluate whether c-MYC depletion also enhanced functional neuronal properties, we performed calcium imaging on neurons transduced with *MYC*-targeting shRNA. Compared to neurons transduced with a scrambled control shRNA, c-MYC-depleted neurons exhibited markedly increased calcium influx in response to KCl stimulation, indicating enhanced excitability (Figure 6E). This functional improvement mirrors the calcium response observed in corrected neurons (Figure 2I), further supporting the role of c-MYC downregulation in rescuing neuronal function.

To pharmacologically inhibit c-MYC, we treated L2HGDH-deficient NPCs with the small-molecule c-MYC inhibitor EN4. This compound targets cysteine 171 of c-MYC within a predicted intrinsically disordered region of the protein, reducing c-MYC and MAX thermal stability, and inhibiting MYC transcriptional activity in cells(42).

Immunoblotting revealed a time-dependent decrease in c-MYC levels following EN4 treatment, which correlated with increased Neurogenin-2 expression (Figure 6F).

Quantification of neurite outgrowth in NPCs subjected to EN4 demonstrated that continuous EN4 treatment during both the NPC and neuronal differentiation stages enhanced total neurite length, nearly to the same degree as correcting the *L2HGDH* allele (Figure 6G). In contrast, EN4 treatment limited to the NPC stage alone had no effect compared to controls, indicating that sustained inhibition of c-MYC is necessary to reverse differentiation defects in this model (Figure 6G). To assess functional improvement, we measured neuronal calcium responses using Fura-2 ratio imaging.

EN4-treated neurons exhibited enhanced intracellular calcium influx in response to KCl stimulation compared to DMSO-treated controls (Figure 6H), indicating that pharmacologic suppression of c-MYC not only improves neuronal morphology but also restores neuronal function in L2HGDH-deficient cells.

Altogether these results demonstrate that depletion or inhibition of c-MYC is sufficient to overcome the effects of L2HGDH deficiency in NPCs, promoting neuronal differentiation and neurite outgrowth to a similar effect as restoring the wild-type *L2HGDH* sequence.

## DISCUSSION

Many IEMs manifesting in the central nervous system cause catastrophic episodes of metabolic decompensation, frequently early in life, leading to acute damage to the brain in the setting of acidosis, hyperammonemia, hypoglycemia and other complications of severe, systemic metabolic dysfunction. In contrast, the cerebral organic acidemias, including L2HGDH deficiency, are often characterized by a later or more insidious onset, progressive neurological dysfunction and macrocephaly(43). The pathophysiological mechanisms of neurological dysfunction in cerebral organic acidemias are not completely understood, and the striking clinical differences from decompensating disorders raise the possibility of mechanisms beyond alterations in intermediary metabolism. The chronic nature of L2HGDH deficiency implies that its effects accumulate over time, possibly due to persistent epigenetic and transcriptional abnormalities induced by L-2HG accumulation(44, 45). The association of L2HGDH deficiency with pediatric tumors, particularly in the brain, also implies dysfunction of terminal differentiation programs in this disease. Our findings support this model by demonstrating that L-2HG inhibits demethylation of a subset of histone marks in neural progenitor cells, induces widespread changes in gene expression and suppresses differentiation into neurons. This results in an overexpression of immature cells and a depletion of excitatory neurons in cortical spheroids.

In our models, L2HGDH deficiency promotes NPC proliferation and self-renewal at the expense of neuronal differentiation. A pivotal effect of L-2HG accumulation is the activation of *MYC* expression. c-MYC has myriad effects in support of cell proliferation, and has been reported to contribute to neural progenitor cell identity(46–48) and suppress neuronal differentiation(49, 50). c-MYC’s involvement in the phenotype of L2HGDH-deficient NPCs is further supported by its functions in glioma, including glioma stem cell maintenance(51, 52). Substantial evidence connects *MYC* specifically to the pathophysiology of IDH1/2-mutant gliomas, with elevated nuclear c-MYC expression in most of these tumors(53). The *MYC* locus has been identified as the most frequently amplified genomic region during glioma malignant progression, with such amplification correlating with increased tumor aggressiveness(54). Furthermore, rs55705857, a noncoding single-nucleotide polymorphism, is associated with a 6-fold increased risk of developing IDH-mutant gliomas with mutations in *IDH1* or *IDH2*(55). This locus lies within a brain-specific enhancer, with the risk allele promoting interaction with the *MYC* locus and driving *MYC* expression(56). Engineering mice to contain the orthologous risk allele is sufficient to increase the penetrance and decrease the latency of IDH1-mutant gliomas(56).

Among the many possible epigenetic effects of elevated L-2HG in *L2HGDH*-deficient cells, we focused on changes predicted to enhance transcription, particularly di- and trimethylation of H3K4, because of the high levels of *MYC* expression. ChIP-seq revealed thousands of genomic sites with altered H3K4me2 and H3K4me3 between unedited and corrected NPCs, including increased abundance in the *MYC* regulatory region. These marks likely resulted from suppression of KDM5 demethylases, because these enzymes are potently inhibited by L-2HG and because treating corrected cells with a KDM5 inhibitor phenocopied the effect of L2HGDH deficiency on these marks.

Despite the widespread changes of H3K4me2 and H3K4me3, genetically or pharmacologically suppressing a single gene – *MYC –* was sufficient to induce levels of neurite formation similar to the effect of editing mutant *L2HGDH* to wild-type. It is possible that other processes beyond H3K4 methylation regulate *MYC* in these models, because KDM5 inhibition had a large effect on H3K4 methylation but more modest effects on *MYC* expression and neurite formation. DNA demethylation(10) and RNA demethylation(34, 57) also involve α-ketoglutarate-dependent dioxygenases and can be blocked by 2HG in some systems. We did not observe global differences in DNA or RNA methylation between unedited and corrected cells, but we cannot rule out localized effects that might impact *MYC*. Nevertheless, our data indicate that high levels of L-2HG promote a chromatin state that favors *MYC* transcription, and it is remarkable that neuronal differentiation could be so effectively stimulated by manipulating this single gene among the thousands of loci altered by L2HGDH deficiency.

Because human L2HGDH deficiency involves progressive, postnatal neurological compromise, there may be opportunities to block disease progression using drugs that interrupt the biological effects of high L-2HG. Along these lines, the chronic, developmental nature of L2HGDH deficiency places the therapeutic focus on long-term management, which perhaps could include therapies aimed at suppressing neural stem cell self-renewal or stimulating neuronal differentiation. We report two different drugs that stimulate neurite formation in L2HGDH-deficient NPCs: EN4, a small molecule that reduces c-MYC stability(42), and CB-839, a glutaminase inhibitor. The mechanism by which CB-839 stimulates neurite formation is through depletion of L-2HG, because glutamine is the predominant source of α-ketoglutarate in these cultured cells. It should be emphasized that this likely not the case in the brain, which supplies the TCA cycle with other fuels, particularly pyruvate. But the conceptual importance of these therapeutic experiments is that the differentiation defect in NPCs can be reversed using multiple different pharmacological approaches. Future studies should explore the efficacy of postnatal therapeutic interventions in preclinical models, particularly in targeting epigenetic and transcriptional dysregulation, to determine whether early treatment can alter the disease course and improve neurological function.

Both L-2HG and D-2HG have been demonstrated to interfere with epigenetic reprogramming by altering the function of α-ketoglutarate-dependent dioxygenases. This would seem to imply that the two enantiomers have largely redundant biological effects. However, in vitro studies have revealed markedly different inhibitory potencies of these molecules on specific dioxygenases, with distinct *Ki* values reported for enzymes such as the TET family of hydroxylases and KDM family of histone demethylases. These biochemical differences indicate that their downstream consequences are not identical, and that the relative sensitivity of epigenetic regulators to each enantiomer could drive divergent biological outcomes(38). In this context, the disparate clinical and developmental phenotypes of diseases associated with high L-2HG or D-2HG are intriguing. L-2HG elevation in the setting of germline *L2HGDH* mutations increases the risk of pediatric tumors in the brain and elsewhere. In adults, D-2HG elevation in the setting of somatic mutations in *IDH1* or *IDH2* also promote tumorigenesis in the brain and some other organs. Yet diseases that cause systemic elevations of D-2HG in childhood have not been reported to have tumors in the brain.

Isolated D-2HGA can be caused by bi-allelic mutations in *D2HGDH* or by monoallelic gain-of-function mutations in *IDH1/2*. These patients have neurodevelopmental disability and, in states of very high D-2HG, cardiomyopathy. Myeloid leukemias were reported in two children, one of whom also had a complex chromosomal rearrangement(58, 59), and other children have had benign tumors(60). It is unknown why these patients seem not to have a high risk of brain tumors despite the clear association between D-2HG and gliomas in adults. This suggests that the impact of L-2HG and D-2HG depends on the lineage and perhaps the developmental stage. Isogenic, patient-derived iPSC models that can be differentiated into different cell types should be useful to decode relationships between dioxygenase expression, epigenetics, lineage specification and the two enantiomers of 2HG.

## METHODS

### Sex as a biological variable

This study involved human-derived induced pluripotent stem cells (iPSCs) and mouse-derived neural progenitor cells (NPCs). The primary iPSC lines were generated from two female patients with L2HGDH deficiency. Matched isogenic controls were derived through CRISPR/Cas9-mediated gene correction. Because these cell lines are clonal and genetically matched apart from the *L2HGDH* mutation, sex-based differences were not a variable in the design or interpretation of experiments. In mouse studies, E15.5 embryos were used without sex determination, as sex cannot be reliably distinguished at this stage of development. While sex was not explicitly considered as a biological variable in this study, the core findings related to L-2HG accumulation and its downstream effects on epigenetic regulation and neuronal differentiation are anticipated to be relevant to both sexes.

### Human fibroblast culture

Human fibroblast lines were derived from skin biopsies or obtained from commercial sources. Punch biopsies of skin for fibroblast culture were obtained from patients following standard clinical diagnostic procedures. All subjects were enrolled in a study (NCT02650622) approved by the Institutional Review Board (IRB) at the University of Texas Southwestern Medical Center (UTSW), and informed consent was obtained from all patients and their families. Fibroblasts were cultured in Dulbecco’s Modified Eagle Medium (DMEM, Sigma D5796) supplemented with 10% fetal bovine serum (FBS), 1% non-essential amino acids (NEAA, GIBCO 11140-050), and 1% penicillin-streptomycin (Pen-Strep). Cultures were incubated at 37°C in a humidified atmosphere containing 5% CO₂.

For routine passaging, when the cells reached 80–90% confluency, they were dissociated using 0.05% trypsin-EDTA and replated at a 1:3 to 1:5 dilution. Fibroblasts were regularly tested for mycoplasma contamination using a PCR-based assay and maintained for no more than 10 passages to prevent cellular senescence.

For experimental assays, fibroblasts were seeded at the desired density and allowed to adhere overnight before treatments or downstream applications. Cells were harvested by washing with cold PBS followed by enzymatic dissociation or direct lysis, depending on the requirements of subsequent analyses.

### D-2HG and L-2HG quantitation

To quantify D-2-hydroxyglutarate (D-2HG) and L-2-hydroxyglutarate (L-2HG), metabolites were extracted from cells grown in 6-cm culture dishes using 80% methanol–water solution. The resulting supernatant was dried in a SpeedVac concentrator, and the dried pellet was resuspended in a diacetyl-L-tartaric anhydride derivatization solution (90 µL of 50 mg/mL diacetyl-L-tartaric anhydride in freshly prepared 80% acetonitrile/20% acetic acid; Acros Organics). To account for sample variability, [U-^13^C]D/L-2HG (Cambridge Isotope Laboratories, 10 ng in 10 µL acetonitrile) was added as an internal standard for unlabeled samples.

The derivatization mixture was sonicated, incubated at 75°C for 30 minutes, then cooled to room temperature before centrifugation to collect the supernatant. The supernatant was again dried in a SpeedVac and reconstituted in 100 µL of 1.5 mmol/L ammonium formate aqueous solution with 10% acetonitrile for liquid chromatography-mass spectrometry (LC-MS) analysis.

Metabolite separation and detection were performed using an AB Sciex 5500 QTRAP LC-MS system (Applied Biosystems SCIEX) equipped with a triple quadrupole/ion trap mass spectrometer and an electrospray ionization (ESI) interface, controlled by AB Sciex Analyst 1.6.1 software. Samples were injected onto a Waters Acquity UPLC HSS T3 column (150 × 2.1 mm, 1.8 µm) for chromatographic separation. The mobile phase consisted of 1.5 mmol/L ammonium formate aqueous solution (pH 3.6, adjusted with formic acid) (Solvent A) and pure acetonitrile (Solvent B). The gradient elution program was as follows:

- 0–12 min: Linear gradient from 1% to 8% B
- 12–15 min: Increased to 99% B
- 15–20 min: Column wash with 99% B
- 20–23 min: Re-equilibration with 1% B

The column was maintained at 35°C, with a flow rate of 0.25 mL/min. Multiple reaction monitoring (MRM) was used to detect 2-hydroxyglutarate-diacetyl tartrate derivatives with the following transitions:

- m/z 363 → 147 (M + 0, CE: -14V)
- m/z 368 → 152 (M + 5, CE: -14V)

### iPSC generation

Pluripotent stem cell experiments were conducted in accordance with guidelines established by the UTSW Stem Cell Research Oversight Committee. To model L-2-hydroxyglutaric aciduria (L-2HGA) in vitro and investigate its impact on neuronal differentiation, induced pluripotent stem cells (iPSCs) were generated from patient fibroblasts using the CytoTune™-iPS 2.0 Sendai Reprogramming Kit (Thermo Fisher Scientific). This kit delivers the Yamanaka reprogramming factors—OCT4, KLF4, SOX2, and c-MYC—via a non-integrating Sendai virus (SeV), allowing for efficient and footprint-free reprogramming.

Patient fibroblasts were cultured in fibroblast medium (DMEM supplemented with 10% fetal bovine serum, 1% nonessential amino acids, 1% penicillin-streptomycin, and 0.1 mM β-mercaptoethanol) until they reached 30–60% confluency. Cells were transduced with the Sendai virus vectors at the recommended multiplicity of infection (MOI) of 5:5:3 (KOS: MYC: KLF4). After 24 hours, the medium was replaced with fresh fibroblast medium, and cells were cultured for an additional six days before passaging onto irradiated mouse embryonic fibroblast (MEF) feeder layers. Seven days post-transduction, cells were transferred to MEF-coated dishes in human iPSC medium (DMEM/F-12, 20% KnockOut™ Serum Replacement, 1% nonessential amino acids, 0.1 mM β-mercaptoethanol, and 10 ng/mL bFGF). Media were replaced daily, and colonies with human embryonic stem cell-like morphology began to emerge within 12–21 days.

For each patient, eight clonal iPSC lines were derived, two of which were further expanded and characterized. Reprogramming efficiency was confirmed through immunofluorescence for pluripotency markers (OCT4, SOX2, NANOG, TRA-1-60, and TRA-1-81). Clearance of Sendai virus was monitored by RT-PCR at passage 10. Only virus-free clones were used for downstream differentiation experiments.

### iPSC CRISPR genome editing

The PX458 cloning with single-step digestion-ligation protocol was used to generate a Cas9-sgRNA expression plasmid containing a GFP marker for subsequent cell sorting. The sgRNA was designed to specifically target the L2HGDH c.829C>T mutation and was cloned into the PX458 vector following single-step digestion-ligation. Plasmid integrity and sgRNA incorporation were verified by Sanger sequencing before use.

Patient-derived iPSCs were maintained in feeder-free conditions and dissociated into single cells using Accutase. The electroporation mixture contained PX458 plasmid (harboring Cas9 and sgRNA) and the ssODN repair template, resuspended in electroporation solution to a final volume of 100 µL per electroporation. Cells were electroporated using the Amaxa Human Stem Cell Nucleofector Kit 1 (Lonza) with a Lonza Nucleofector device under program B-016, following manufacturer guidelines for optimal editing efficiency. Immediately after nucleofection, cells were plated in mTeSR1 medium supplemented with 10 µM ROCK inhibitor to enhance survival.

Forty-eight hours post-electroporation, GFP^+^ cells were isolated via fluorescence-activated cell sorting (FACS) to enrich for successfully transfected cells. Sorted cells were seeded at clonal density and expanded in mTeSR1 medium on Matrigel-coated plates. Individual iPSC colonies were manually picked and expanded for genotyping. Restriction fragment length polymorphism (RFLP) analysis was performed using BtgI, which selectively digests the corrected alleles while leaving unedited alleles intact.

Clones that exhibited the expected digestion pattern were further validated by Sanger sequencing to confirm successful HDR-mediated correction of L2HGDH.

### Neural progenitor cell (NPC) generation

To differentiate iPSCs into neural progenitor cells (NPCs), we followed a dual-SMAD inhibition neural specification protocol as described(26). This protocol efficiently drives neuroectodermal differentiation while suppressing alternative mesodermal and endodermal fates, enabling the robust generation of NPCs. iPSCs were cultured in feeder-free conditions and maintained in mTeSR1 medium on Matrigel-coated plates until reaching ∼70–80% confluency. To initiate neural differentiation, cells were transitioned to N2/B27 neural induction medium, supplemented with 0.1 µM LDN193189 and 10 µM SB431542, small-molecule inhibitors of BMP and TGF-β signaling, respectively. Neural rosettes, characteristic of neuroepithelial differentiation, began to emerge within 7–10 days. On day 14, neural rosettes were mechanically isolated and transferred to Poly-L-Ornithine/Laminin-coated plates in NPC expansion medium containing N2, B27, and 20 ng/mL FGF2. Cultures were maintained with media changes every other day, and cells were routinely passaged using Accutase. NPC identity was validated by immunocytochemistry for NESTIN and SOX2, confirming their neural progenitor state.

### Cortical spheroid generation and measurement

Cortical spheroids were generated from H9 control NPCs, unedited NPCs, and corrected NPCs using a previously established differentiation protocol(27). NPCs were cultured in low-attachment plates and transitioned to cortical spheroid differentiation medium, which was supplemented with dual-SMAD inhibitors (LDN193189 and SB431542) during early neural induction. After 14 days, spheroids were transferred to neural maturation medium containing BDNF and NT3, supporting the progressive differentiation of neurons and glial cells. After 30 days of differentiation, spheroids were imaged using a Zeiss SteREO Discovery V.8 stereomicroscope under brightfield conditions. To quantify spheroid size, images were processed in ImageJ, where spheroid surface area was measured and compared across groups (unedited, corrected, and control NPC-derived spheroids).

### Clonogenicity assay for NPCs

To assess NPC self-renewal capacity, we performed a clonogenicity assay using a modified Neural-Colony Forming Cell Assay (N-CFCA) protocol(28). NPCs were dissociated into a single-cell suspension using Accutase and passed through a 40-μm cell strainer to remove cell clumps. The plated cultures were transferred to a humidified incubator (37°C, 5% CO₂) and were replenished with fresh medium containing growth factors every 7 days for a total of 21 days, allowing cells to reach their full proliferative potential. On day 21, colonies were imaged using stereoscopic and phase-contrast microscopy. To quantify sphere-forming frequency, colonies were counted using ImageJ.

### Neuronal differentiation and neurite imaging and quantification

NPCs were dissociated into a single-cell suspension using Accutase (4 min at room temperature), followed by neutralization with NPC medium and centrifugation at 300g for 5 min. The pellet was resuspended in a small volume of neuronal differentiation medium, and cells were counted and plated onto poly-L-ornithine (PLO)/laminin-coated plates at a density of 200,000 cells per well (6-well plate). Cells were maintained in neuronal differentiation medium, consisting of DMEM/F12 supplemented with 1× N2 supplement, 1× B27 supplement (without retinoic acid), 20 ng/mL BDNF (Peprotech), 20 ng/mL GDNF (Peprotech), 1 mM dibutyryl-cAMP (Sigma), 200 nM ascorbic acid (Sigma), and 1 µg/mL laminin (Gibco). The medium was changed every three days, and cultures were maintained for 2–6 weeks to allow for neuronal maturation.

On day 14, differentiated neurons were immunostained for MAP2 and β-III-Tubulin, and confocal images were acquired using a Spinning Disk Confocal Nikon CSU-W1 with SoRa. Each image covered an area of 6325.48 × 6325.48 µm, generated by stitching 9 × 9 tiled images. Neurite outgrowth was analyzed using the SNT (Simple Neurite Tracer) plugin in Fiji ImageJ, which was used to trace and quantify the total neurite lengths of individual neurons.

### Isolation and culture of mouse neural progenitor cells (NPCs)

Neural progenitor cells were isolated from embryonic day 15.5 (E15.5) mouse embryos as previously described(61) with modifications. Embryos were collected in ice-cold 1× PBS, and the dorsal telencephalon was microdissected under a stereoscope. Meninges were carefully removed using fine forceps. Dissected tissue was transferred to a 15-mL conical tube containing 1 mL of prewarmed enzyme solution composed of dissociation medium supplemented with papain (20 U/mL; Sigma) and 6.4 mg/mL cysteine. Tissue was digested at 37 °C for 20 minutes, gently mixed, and incubated for an additional 20 minutes after the addition of a second 1 mL of enzyme solution.

Following enzymatic digestion, tissue was washed twice with light inhibitory solution (LI), followed by a brief 2-minute incubation in heavy inhibitory solution (HI) at 37 °C. The tissue was then rinsed in NEP complete medium and gently dissociated into a single-cell suspension by trituration with a fire-polished pipette in 0.5–1.0 mL NEP complete medium. Cells were counted using a hemocytometer and plated at a density of 1×10⁶ cells per 6-cm nontreated polystyrene dish.

NEP complete medium was prepared by supplementing NEP basal medium with 100 ng/mL epidermal growth factor (EGF) and 10 ng/mL basic fibroblast growth factor (bFGF). NEP basal medium consisted of Neurobasal medium (Invitrogen 21103049) supplemented with 1× GlutaMAX (Gibco 35050-061), 1× normocin (Fisher NC9273499), 1× B27-RA (Gibco 12587-010), and 1× N2 supplement (Gibco 17502-048). Cultures were maintained at 37 °C in a humidified incubator with 5% CO₂. Neurosphere formation was typically observed within 3–5 days. Medium was replaced every three days by allowing spheres to settle by gravity, aspirating spent medium, and replenishing with fresh NEP complete medium.

For differentiation, neurospheres were plated onto poly-D-lysine/laminin-coated eight-well chamber slides and maintained in NEP basal medium supplemented with 2% heat-inactivated horse serum (Gibco 26050070) for 2–3 days, with daily medium changes.

For neurite analysis, cultures were fixed in 4% paraformaldehyde and immunostained for neuronal markers.

### Ca²⁺ imaging

Ca²⁺ imaging experiments were performed on neuronal cultures at day 47 following previously established protocols (Zhang et al., 2006). Briefly, neurons were loaded with 5 µM Fura-2 AM (Molecular Probes) and incubated for 30 min at 37°C in Tyrode solution to allow for dye uptake and de-esterification. Coverslips containing neurons were then mounted onto a recording/perfusion chamber (RC-26G, Warner Instruments) positioned on the movable stage of an Olympus IX-70 inverted microscope.

Neuronal activity was assessed by stimulating neurons with a KCl-depolarizing solution (45 mM KCl in Tyrode solution). Ca²⁺ responses were measured via intermittent excitation at 340 nm and 380 nm UV light using a DeltaRAM illuminator (PTI) in combination with a Fura-2 dichroic filter cube (Chroma Technologies) and a 60× UV-grade oil-immersion objective (Olympus).

The emitted fluorescence signal was captured using an IC-300 camera (PTI), and images were digitized and analyzed using ImageMaster Pro software (PTI). Baseline Ca²⁺ levels were recorded before KCl stimulation, and images were acquired at 340 nm and 380 nm excitation wavelengths every 5 s, with the peak fluorescence value measured post-stimulation. Background fluorescence was determined according to manufacturer guidelines (PTI) and subtracted from the signal to ensure accurate quantification of Ca²⁺ transients.

### Histone extraction

Histone proteins were extracted from cultured cells following a standard acid extraction protocol to preserve post-translational modifications. Cells were grown in 15-cm culture dishes until reaching ∼70% confluency and then washed twice with ice-cold PBS supplemented with 5 mM sodium butyrate to prevent histone deacetylation. Cells were collected by scraping in 1 mL of PBS with sodium butyrate, followed by centrifugation at 1,000g for 5 min at 4°C to pellet the cells. The supernatant was discarded, and the pellet was resuspended in Triton Extraction Buffer (TEB) at a density of 1 × 10⁷ cells/mL. TEB consisted of PBS supplemented with 0.5% Triton X-100 (v/v), 2 mM phenylmethylsulfonyl fluoride (PMSF), and 0.02% (w/v) sodium azide (NaN₃). Cells were lysed on ice for 10 min with gentle rocking, then centrifuged at 1,000g for 10 min at 4°C to pellet nuclei. The nuclear pellet was washed with 0.5× volume of TEB, followed by another centrifugation at 1,000g for 10 min at 4°C. The pellet was then resuspended in 0.2N HCl at a final concentration of 4 × 10⁷ cells/mL and incubated overnight at 4°C on a rotating platform to extract histones. The following day, samples were centrifuged at 1,000g for 10 min at 4°C, and the acid-extracted histones in the supernatant were collected. Protein concentration was determined using the Bradford assay, and aliquots were stored at -80°C for long-term use. For Western blot analysis, 2 µg of histone extract was loaded onto a polyacrylamide gel, and samples were denatured before electrophoresis.

### DNA dot blot analysis

To assess global DNA methylation levels, DNA dot blot analysis was performed using a Bio-Dot Microfiltration Apparatus (Bio-Rad). Genomic DNA was isolated using a standard DNA purification protocol, and its concentration was measured using a Qubit fluorometer. DNA was then fragmented via sonication using a Covaris ME220 instrument, generating fragments between 200–500 bp. DNA quality and fragmentation efficiency were assessed using agarose gel electrophoresis or TapeStation analysis.

Fragmented DNA was diluted to 50 ng/µL in nuclease-free water, and 100 µL of DNA was mixed with an equal volume of 2× DNA denaturing buffer (200 mM NaOH, 20 mM EDTA). The samples were incubated at 95°C for 10 min to denature the DNA, followed by immediate chilling on ice for 5 min. After denaturation, 200 µL of 20× SSC buffer was added to each sample to neutralize the pH, and the final volume was adjusted to 500 µL with nuclease-free water. A six-point serial dilution was prepared from the denatured DNA.

A positively charged nylon membrane (Zeta-Probe, Bio-Rad) was pre-wetted in 2× SSC buffer for 10 min. Using the Bio-Dot apparatus, 100 µL of each DNA dilution was applied to individual wells and allowed to filter through under gentle vacuum. After blotting, the membrane was rinsed once with 500 µL of 0.4 M NaOH (for Zeta-Probe membranes) or 2× SSC (for nitrocellulose membranes) to ensure proper DNA binding. The membrane was then UV cross-linked at 1200 J/m² using a Stratalinker UV cross-linker (Stratagene) to fix the DNA to the membrane.

Membranes were blocked in TBST containing 5% nonfat dry milk for 1 hour at room temperature, followed by incubation overnight at 4°C with primary antibodies targeting specific DNA modifications. Following three washes in 1× TBST (5 min each), membranes were incubated with an HRP-conjugated secondary antibody for 1 hour at room temperature. The membrane was washed again three times for 5 min each and then incubated with ECL reagent for 1 min at room temperature. Chemiluminescent signals were captured using a GE Imager.

### RNA dot blot analysis

RNA dot blot analysis was performed using a Bio-Dot Microfiltration Apparatus (Bio-Rad) to assess global RNA modifications. Total RNA was extracted using a standard RNA isolation protocol, and RNA integrity was assessed using ScreenTape analysis (Agilent) to confirm the presence of distinct 28S and 18S rRNA bands, ensuring sample integrity. RNA concentrations were measured using a Qubit fluorometer, and samples were diluted 20-fold in nuclease-free water before further processing.

To prepare samples for blotting, RNA was diluted to 160 ng/µL in a final volume of 50 µL, followed by denaturation with 4× RNA denaturing buffer (16.4M formamide, 2.8M formaldehyde, 26.6mM MOPS buffer) at 65°C for 5 min. The reaction was immediately chilled on ice for 5 min before adding 66.5 µL of 20× SSC buffer, followed by 67 µL of nuclease-free water to achieve a final RNA concentration of 40 ng/µL in 200 µL. A six-point, two-fold dilution series was prepared by sequentially diluting RNA samples in nuclease-free water.

A positively charged nylon membrane (Zeta-Probe, Bio-Rad) was pre-wet by slowly submerging it at a 45° angle into distilled water, followed by equilibration in 2× SSC buffer. The Bio-Dot apparatus was assembled according to the manufacturer’s instructions, and all wells were rinsed with 500 µL of 2× SSC buffer before sample loading. The RNA was allowed to bind the membrane under gentle vacuum pressure, ensuring that the membrane was mostly dry before proceeding. Following RNA immobilization, the membrane was rinsed with 500 µL of 2× SSC with 0.1% SDS, removed from the apparatus, and further washed in 2× SSC buffer.

The membrane was UV cross-linked at 1200 J/m² using a Stratalinker UV cross-linker (Stratagene) to covalently fix RNA. Membranes were then blocked and probed with primary antibodies against RNA modifications. Following overnight incubation at 4°C, membranes were washed three times in 1× TBST, incubated with HRP-conjugated secondary antibodies, and visualized using enhanced chemiluminescence (ECL) imaging on a GE Imager.

### Human pluripotent stem cell culture

The human embryonic stem cell (ESC) line H9 and human induced pluripotent stem cell (hiPSC) lines (WiCell Research Institute, Madison, WI) were maintained on irradiated mouse embryonic fibroblast (MEF) feeder layers (GlobalStem). Cells were passaged enzymatically by treating with 1 mg/ml Collagenase Type IV (STEMCELL Technologies) for 2 minutes, followed by mechanical lifting with a cell scraper (Falcon).

Feeder-supported human pluripotent stem cells (PSCs) were cultured in hPSM medium consisting of DMEM/F12 (Invitrogen) supplemented with 20% KnockOut Serum Replacement (KSR; Invitrogen), 1% non-essential amino acids (Invitrogen), 0.1 mM β-mercaptoethanol (Invitrogen), and 10 ng/ml FGF2 (Peprotech). Feeder-free PSCs were cultured on Matrigel (1:25 dilution; BD Biosciences) in mTeSR1 medium (STEMCELL Technologies).

All tissue culture media were filtered using 0.22 μm low protein-binding filters (Millipore). Cells were maintained at 37°C under 5% CO2 and atmospheric oxygen conditions.

### Extract preparation and immunoblot assays

Whole-cell lysates were prepared by lysing cells in M-PER Mammalian Protein Extraction Reagent (Thermo) supplemented with 20 mM NaF, 1 mM Na3VO4, 4 μg/ml aprotinin, 4 μg/ml leupeptin, 4 μg/ml pepstatin, and 1 mM DTT.

Nuclear and cytoplasmic fractions were isolated using a two-step protocol. Cells were first resuspended in Buffer A [10 mM HEPES (pH 7.9), 10 mM KCl, 0.1 mM EDTA, 0.4% NP40, 20 mM NaF, 1 mM Na3VO4, 4 μg/ml aprotinin, 4 μg/ml leupeptin, 4 μg/ml pepstatin, and 1 mM DTT] to extract cytoplasmic proteins. The nuclear pellet was resuspended in Buffer B [20 mM HEPES (pH 7.9), 400 mM NaCl, 1 mM EDTA, 10% glycerol, 20 mM NaF, 1 mM Na3VO4, 4 μg/ml aprotinin, 4 μg/ml leupeptin, 4 μg/ml pepstatin, and 1 mM DTT] and vortexed for 2 hours to extract nuclear proteins.

Western blotting was performed using standard protocols, and protein concentrations of cell extracts were measured by the Bradford assay.

### Targeted metabolomics

Metabolites were extracted by rinsing cells twice with ice-cold saline, followed by quenching with 80% cold methanol. Cells were incubated at -80°C for at least 20 minutes before being scraped. Tissue samples were thawed and homogenized in cold 80% methanol using plastic pestles (Thermo Fisher Scientific, 12141364).

Samples underwent three freeze-thaw cycles, alternating between liquid nitrogen and a 37°C water bath. After the final thaw, samples were vortexed for 1 minute and centrifuged at 20,160 × g for 15 minutes at 4°C. The supernatants were transferred to fresh Eppendorf tubes and dried overnight in a SpeedVac concentrator.

For analysis on a Q-Exactive mass spectrometer, dried samples were reconstituted in 80% acetonitrile and centrifuged at 20,160 × g at 4°C to remove insoluble material.

Metabolites were separated chromatographically using a Vanquish UHPLC system equipped with a ZIC-pHILIC column (Millipore-Sigma, Burlington, MA), as described previously(62–64).

Extracted ion chromatograms (XICs) were generated with a mass tolerance of 5 ppm and integrated for relative quantitation. Analyte identities were confirmed by comparison with purified standards and product ion spectra.

### Stable isotope tracing

For ^13^C-glucose or ^13^C-glutamine tracing, cells were cultured in DMEM/F12 base medium supplemented with 17.5 mM [U-^13^C]glucose (Cambridge Isotope Laboratories, CLM-481) or 2.5 mM [U-^13^C]glutamine (Cambridge Isotope Laboratories, CLM-1822). Metabolite extraction was performed as described in the targeted metabolomics section.

Natural isotope abundances were corrected using a customized R script, adapted from the AccuCor algorithm (65), and made publicly available at the following GitHub repository: https://github.com/wencgu/nac.

### Immunofluorescence and confocal microscopy

Cells were seeded onto coverslips and fixed the following day with freshly prepared 4% paraformaldehyde (PFA) in PBS for 15 minutes. After fixation, cells were permeabilized with 0.1% (v/v) Triton X-100 in PBS at room temperature for 10 minutes.

Blocking was performed in PBS containing 1% BSA for at least 30 minutes at room temperature. Cells were then incubated with primary antibodies for 1 hour at room temperature. Following primary antibody incubation, cells were washed three times with PBS (5 minutes each) and incubated with Alexa Fluor-conjugated secondary antibodies (Alexa Fluor 488 and 555, Invitrogen) for 3 hours at room temperature in the dark.

Coverslips were washed three times with PBS (5 minutes each) and once with Milli-Q water, then mounted onto slides using ProLong™ Glass Antifade Mountant (P36935, Invitrogen) and left to cure overnight in the dark.

Images were acquired using a Zeiss LSM 880 Confocal Laser Scanning Microscope with Z-stacks captured. Image processing and analysis were performed using Fiji (ImageJ).

### RNA sequencing (RNA-seq)

Total RNA was extracted using TRIzol™ Reagent (Thermo Fisher Scientific, 15596018) followed by purification with the RNeasy Mini Kit (Qiagen, 74106). RNA concentrations were measured using a Qubit™ Fluorometer and the Qubit RNA High Sensitivity Kit (Invitrogen, Q32852).

RNA-seq libraries were prepared using the NEBNext Ultra II Directional RNA Library Prep Kit with the NEBNext Poly(A) mRNA Magnetic Isolation Module (New England Biolabs, E7490L, E7760L) according to the manufacturer’s instructions. Library indexing was performed using the NEBNext Multiplex Oligos for Illumina (E7730L, E7335L, E7500L).

Sequencing reads were aligned to the hg19 reference genome using STAR v.2.5.2b(66) with the following parameters:

--runThreadN 28

--outSAMtype BAM SortedByCoordinate

--outFilterMultimapNmax 1

--outWigStrand Unstranded

--quantMode TranscriptomeSAM

The resulting BAM files were converted to BED format using the bamtobed command in BEDtools v.2.29.2 (https://bedtools.readthedocs.io/en/latest/). BED files were normalized and converted to wiggle format using a custom Python script. Wiggle files were subsequently converted to bigWig format with the wigToBigWig tool, applying the - clip parameter.

Gene-level read counts were obtained using HTSeq(67) with the parameter -s no. To ensure uniformity, one additional read count was added to each gene for each independent sample prior to downstream analysis.

Differentially expressed genes were identified using DESeq2(68), with the following thresholds applied:

- Fold change ≥ 1.5
- FDR-adjusted P value ≤ 0.05

### Chromatin immunoprecipitation sequencing (ChIP-seq)

ChIP-seq was performed as previously described(69). Briefly, crosslinked chromatin was sonicated in RIPA 0 buffer (10 mM Tris-HCl, 1 mM EDTA, 0.1% sodium deoxycholate, 0.1% SDS, 1% Triton X-100, 0.25% Sarkosyl, pH 8.0) to achieve fragment sizes between 200 and 500 bp. The chromatin was adjusted to a final concentration of 150 mM NaCl and incubated overnight at 4°C with the appropriate antibodies.

Following immunoprecipitation, chromatin was washed, and ChIP DNA was purified. ChIP-seq libraries were prepared using the NEBNext Ultra II DNA Library Prep Kit (New England Biolabs) according to the manufacturer’s protocol and sequenced on an Illumina NextSeq 500 platform using the 75 bp high-output sequencing kit.

Raw ChIP-seq reads were aligned to the human hg19 genome assembly using Bowtie2 (70) with default parameters. Only uniquely mapped reads were retained for further analysis. Peak calling was performed using MACS with the --nomodel parameter(71).

### Single-cell RNA sequencing

Human cortical spheroids (hCSs) were generated following the Pasca laboratory protocol and harvested at day 45 for single-cell dissociation(27). Organoids were enzymatically dissociated using Accutase (Innovative Cell Technologies) at 37°C for 15 minutes, followed by gentle mechanical trituration. Single-cell suspensions were filtered through a 40 μm cell strainer, and viability was confirmed using trypan blue exclusion.

Cells were immediately loaded onto a Chromium Controller (10x Genomics) using the Chromium Next GEM Single Cell 3’ kit v3.1 (10x Genomics), targeting 5,000–10,000 cells per sample. Reverse transcription, cDNA amplification, and library preparation were performed according to the manufacturer’s protocol. Libraries were sequenced on an Illumina NovaSeq 6000 using paired-end 150 bp reads.

Raw FASTQ files were processed using Cell Ranger (v6.1.2) with default settings and aligned to the human genome (GRCh38). Output gene-cell matrices were filtered to retain only high-quality cells (e.g., cells with >300 and <7500 detected genes and <15% mitochondrial transcripts) using Seurat (v5.0.1) in R.

Samples were individually normalized using SCTransform and integrated using canonical correlation analysis with Seurat’s FindIntegrationAnchors() and IntegrateData() functions (normalization.method = “SCT”). The integrated object was subjected to dimensionality reduction by principal component analysis (PCA). Uniform Manifold Approximation and Projection (UMAP) was used to embed cells in 2D space using the first 12 principal components.

Clusters were identified using the Louvain algorithm implemented in Seurat’s FindClusters() function at a resolution of 0.5. The RunUMAP() function was used for visualization. Marker genes for each cluster were identified using FindAllMarkers() with parameters only.pos = TRUE, min.pct = 0.25, and logfc.threshold = 0.25.

Cell type identities were assigned by integrating canonical marker expression profiles, literature-based reference signatures, and automatic label transfer. Initial annotations were performed using SingleR with the Human Primary Cell Atlas as reference. These were further refined manually by examining expression of key lineage markers (e.g., *SOX2*, *PAX6*, *NEUROG2*, *VIM*, *STMN2*, *GAD1*, *SLC17A7*, *GFAP*, *NFIA*, *DCX*, *TUBB3*), with final labels assigned to 21 distinct cell populations representing radial glia, intermediate progenitors, immature neurons, excitatory and inhibitory neurons, oligodendrocyte precursor cells (OPCs), and astrocyte precursors.

To compare *MYC* expression between corrected and unedited hCSs, cells were grouped by cell_type and condition, and average expression values were calculated using FetchData() and group_by(). For each cell type, statistical significance was assessed using the Wilcoxon rank-sum test, and false discovery rate correction was applied using the Benjamini-Hochberg method. Differential expression was visualized using violin plots (VlnPlot) and boxplots with ggplot2.

Proportional representation of each annotated cell type was calculated per sample. The relative frequency of “Immature neuron” subtypes was compared between corrected and unedited conditions using summary statistics and bar plots. Significance of differences in cell-type frequencies was evaluated using non-parametric statistical tests when appropriate.

### Statistics

All statistical analyses were performed using R (version 4.3.1). Data are presented as mean ± SEM unless otherwise noted. For comparisons between two groups, statistical significance was determined using unpaired, two-tailed Student’s t-tests assuming equal variance, unless stated otherwise. For comparisons involving three or more groups, one-way or two-way ANOVA was used as appropriate, followed by Tukey’s HSD or Dunn’s post hoc test to adjust for multiple comparisons. For non-normally distributed datasets, normality was assessed using the Shapiro–Wilk test, and non-parametric tests (e.g., Kruskal–Wallis with Dunn’s correction) were applied where applicable. Statistical details, including sample sizes, exact tests used, and multiple testing corrections, are provided in the corresponding figure legends. A p-value < 0.05 was considered statistically significant.

### Study approval

All studies involving human subjects were approved by the Institutional Review Board (IRB) at the University of Texas Southwestern Medical Center (UTSW). Written informed consent was obtained from all participants or their legal guardians prior to enrollment, including consent for the collection and use of skin biopsies to derive fibroblast and iPSC lines. All procedures involving animals were approved by the UTSW Institutional Animal Care and Use Committee (IACUC).

### Data availability

All sequencing data generated in this study have been deposited in the Gene Expression Omnibus (GEO) under accession numbers GSE295010 (RNA-seq) and GSE295011 (ChIP-seq). Additional raw and processed data supporting the findings of this study are available from the corresponding author upon reasonable request.

## SUPPLEMENTAL MATERIAL

Supplemental material includes supplemental methods, five figures, and eight tables.

## AUTHOR CONTRIBUTIONS

W.G. designed the study, conducted experiments, analyzed results, and wrote the manuscript. X.W. contributed to ChIP-seq experiments. A.S., B.F., Z.W., J.S., L.G.Z., F.C., and T.P.M. contributed to metabolomics experiments. L.C., Y.X., Y.Z., and A.K.W. assisted with sequencing data analysis. A.T., J.F., A.R., H.T., and B.A. provided support with cell culture experiments. H.Z. and I.B. carried out Ca²⁺ imaging experiments and data analysis. S.S. contributed to mouse experiments. S.K.M. provided guidance on the design and interpretation of epigenetic analyses. R.J.D. provided funding, designed the study, analyzed results, and wrote the manuscript.

## ACKNOWLEDGEMENTS

We thank members of the DeBerardinis laboratory and Prashant Mishra for helpful discussions. We thank Hongli Chen, Chunxiao Pan, Danny Vu, Min Ni, K. Celeste Oaxaca, Chendong Yang, Kailong Li, Minghao Li, Yuemeng Jia, Zhimin Gu, Yi Xiao, Chao Xing, Misty Martin-Sandoval, Bookyung Ko, Dennis Dumesnil, Aparna Rao, Varun Sondhi, Tracy Rosales, Divya Bezwada, Sherwin Kelekar, Karla Cano Hernandez and Momoko Watanabe for technical assistance and experimental advice. We are also grateful to the CRI Metabolomics, Sequencing, Moody Flow Cytometry, and Mouse Genome Engineering Core Facilities, as well as the Quantitative Light Microscopy Core at UTSW, for support with data acquisition.

This article is subject to HHMI’s Open Access to Publications policy. HHMI lab heads have previously granted a nonexclusive CC BY 4.0 license to the public and a sublicensable license to HHMI in their research articles. Pursuant to those licenses, the author-accepted manuscript of this article can be made freely available under a CC BY4.0 license immediately upon publication. R.J.D is supported by the Howard Hughes Medical Institute Investigator Program, grants from the National Institutes of Health (NIH) (R35CA220449, P50CA196516, P50CA070907), the Cancer Prevention Research Institute of Texas (CPRIT) (RP220337), the Moody Foundation (Robert L. Moody, Sr. Faculty Scholar Award) and the Eugene McDermott Endowment for the Study of Human Growth and Development. L.G.Z, T.P.M and the CRI Metabolomics Facility are supported by CPRIT (RP240494). S.K.M is supported by NIH (R01CA289260, R01CA258586, P50CA165962, U19CA264504) and CPRIT (RR190034, RP230344, RP2400489, RP250278). S.K.M receives support from Servier Pharmaceuticals and is a co-founder of Gliomet. S.S. is supported by NIH grant R01CA200653, and L.C. is supported by NIH grant R01CA285336. Y.X. is supported by NIH grant K99CA277576 and grant HFSP LT0018/2022-L.

## REFERENCES

1. Van Schaftingen E, Rzem R, Veiga-da-Cunha M. L: -2-Hydroxyglutaric aciduria, a disorder of metabolite repair. J Inherit Metab Dis. 2009;32(2):135–142.

2. Rzem R, et al. A gene encoding a putative FAD-dependent l-2-hydroxyglutarate dehydrogenase is mutated in l-2-hydroxyglutaric aciduria. PNAS. 2004;101(48):16849– 16854.

3. Kranendijk M, et al. Progress in understanding 2-hydroxyglutaric acidurias. Journal of Inherited Metabolic Disease. 2012;35(4):571–587.

4. Moroni I, et al. L-2-hydroxyglutaric aciduria and brain malignant tumors: a predisposing condition? Neurology. 2004;62(10):1882–1884.

5. Rogers RE, et al. Wilms Tumor in a Child with L-2-hydroxyglutaric Aciduria. Pediatr Dev Pathol. 2010;13(5):408–411.

6. Yan H, et al. IDH1 and IDH2 Mutations in Gliomas. New England Journal of Medicine. 2009;360(8):765–773.

7. Mardis ER, et al. Recurring Mutations Found by Sequencing an Acute Myeloid Leukemia Genome. New England Journal of Medicine. 2009;361(11):1058–1066.

8. Dang L, et al. Cancer-associated IDH1 mutations produce 2-hydroxyglutarate. Nature. 2009;462(7274):739–744.

9. Ward PS, et al. The common feature of leukemia-associated IDH1 and IDH2 mutations is a neomorphic enzyme activity converting alpha-ketoglutarate to 2-hydroxyglutarate. Cancer Cell. 2010;17(3):225–234.

10. Xu W, et al. Oncometabolite 2-hydroxyglutarate is a competitive inhibitor of α-ketoglutarate-dependent dioxygenases. Cancer Cell. 2011;19(1):17–30.

11. Chowdhury R, et al. The oncometabolite 2-hydroxyglutarate inhibits histone lysine demethylases. EMBO reports. 2011;12(5):463–469.

12. Figueroa ME, et al. Leukemic IDH1 and IDH2 Mutations Result in a Hypermethylation Phenotype, Disrupt TET2 Function, and Impair Hematopoietic Differentiation. Cancer Cell. 2010;18(6):553–567.

13. Shim E-H, et al. l-2-Hydroxyglutarate: An Epigenetic Modifier and Putative Oncometabolite in Renal Cancer. Cancer Discovery. 2014;4(11):1290–1298.

14. Kundu A, et al. L-2-Hydroxyglutarate remodeling of the epigenome and epitranscriptome creates a metabolic vulnerability in kidney cancer models. J Clin Invest. 2024;134(13). 10.1172/JCI171294.

15. Wise DR, et al. Hypoxia promotes isocitrate dehydrogenase-dependent carboxylation of α-ketoglutarate to citrate to support cell growth and viability. Proceedings of the National Academy of Sciences. 2011;108(49):19611–19616.

16. Oldham WM, et al. Hypoxia-Mediated Increases in l-2-hydroxyglutarate Coordinate the Metabolic Response to Reductive Stress. Cell Metabolism. 2015;22(2):291–303.

17. Intlekofer AM, et al. Hypoxia Induces Production of L-2-Hydroxyglutarate. Cell Metabolism. 2015;22(2):304–311.

18. Intlekofer AM, et al. L -2-Hydroxyglutarate production arises from noncanonical enzyme function at acidic pH. Nature Chemical Biology. 2017;13(5):494–500.

19. Mullen AR, et al. Oxidation of Alpha-Ketoglutarate Is Required for Reductive Carboxylation in Cancer Cells with Mitochondrial Defects. Cell Reports. 2014;7(5):1679– 1690.

20. Ni M, et al. Functional Assessment of Lipoyltransferase-1 Deficiency in Cells, Mice, and Humans. Cell Reports. 2019;27(5):1376–1386.e6.

21. Zhao J, et al. Metabolic remodelling during early mouse embryo development. Nat Metab. 2021;3(10):1372–1384.

22. Li H, et al. Drosophila larvae synthesize the putative oncometabolite L-2-hydroxyglutarate during normal developmental growth. PNAS. 2017;114(6):1353–1358.

23. Li F, et al. Glial Metabolic Rewiring Promotes Axon Regeneration and Functional Recovery in the Central Nervous System. Cell Metabolism. 2020;32(5):767–785.e7.

24. Takahashi K, et al. Induction of Pluripotent Stem Cells from Adult Human Fibroblasts by Defined Factors. Cell. 2007;131(5):861–872.

25. Kwart D, et al. Precise and efficient scarless genome editing in stem cells using CORRECT. Nature Protocols. 2017;12(2):329–354.

26. Topol A, Tran NN, Brennand KJ. A guide to generating and using hiPSC derived NPCs for the study of neurological diseases. J Vis Exp. 2015;(96):e52495.

27. Sloan SA, et al. Generation and assembly of human brain region–specific three-dimensional cultures. Nature Protocols. 2018;13(9):2062–2085.

28. Azari H, et al. Neural-Colony Forming Cell Assay: An Assay To Discriminate Bona Fide Neural Stem Cells from Neural Progenitor Cells. JoVE (Journal of Visualized Experiments). 2011;(49):e2639.

29. Brennand KJ, et al. Modelling schizophrenia using human induced pluripotent stem cells. Nature. 2011;473(7346):221–225.

30. Zhang H, et al. Association of CaV1.3 L-type calcium channels with Shank. J Neurosci. 2005;25(5):1037–1049.

31. Subramanian A, et al. Gene set enrichment analysis: A knowledge-based approach for interpreting genome-wide expression profiles. PNAS. 2005;102(43):15545–15550.

32. Ma Q, Kintner C, Anderson DJ. Identification of neurogenin, a Vertebrate Neuronal Determination Gene. Cell. 1996;87(1):43–52.

33. Brinkley G, et al. Teleological role of L-2-hydroxyglutarate dehydrogenase in the kidney. Disease Models & Mechanisms. 2020;13(11). 10.1242/dmm.045898.

34. Zheng G, et al. ALKBH5 Is a Mammalian RNA Demethylase that Impacts RNA Metabolism and Mouse Fertility. Molecular Cell. 2013;49(1):18–29.

35. Jia G, et al. N6-Methyladenosine in nuclear RNA is a major substrate of the obesity-associated FTO. Nat Chem Biol. 2011;7(12):885–887.

36. Laukka T, et al. Cancer-associated 2-oxoglutarate analogues modify histone methylation by inhibiting histone lysine demethylases. Journal of Molecular Biology. 2018;430(18, Part B):3081–3092.

37. Gunn K, et al. (R)-2-Hydroxyglutarate Inhibits KDM5 Histone Lysine Demethylases to Drive Transformation in IDH-Mutant Cancers. Cancer Discovery. 2023;13(6):1478–1497.

38. Koivunen P, et al. Transformation by the (R)-enantiomer of 2-hydroxyglutarate linked to EGLN activation. Nature. 2012;483(7390):484–488.

39. Liu X, et al. Distinct features of H3K4me3 and H3K27me3 chromatin domains in pre-implantation embryos. Nature. 2016;537(7621):558–562.

40. Klose RJ, et al. The Retinoblastoma Binding Protein RBP2 Is an H3K4 Demethylase. Cell. 2007;128(5):889–900.

41. Horton JR, et al. Structural Basis for KDM5A Histone Lysine Demethylase Inhibition by Diverse Compounds. Cell Chemical Biology. 2016;23(7):769–781.

42. Boike L, et al. Discovery of a Functional Covalent Ligand Targeting an Intrinsically Disordered Cysteine within MYC. Cell Chemical Biology. 2021;28(1):4–13.e17.

43. Wajner M. Neurological manifestations of organic acidurias. Nat Rev Neurol. 2019;15(5):253–271.

44. Kusi M, et al. 2-Hydroxyglutarate destabilizes chromatin regulatory landscape and lineage fidelity to promote cellular heterogeneity. Cell Reports. 2022;38(2):110220.

45. Ahmed S, et al. L-2-hydroxyglutaric aciduria – review of literature and case series. Annals of Medicine and Surgery. 2023;85(4):712.

46. Eilers M, Eisenman RN. Myc’s broad reach. Genes Dev. 2008;22(20):2755–2766.

47. Kerosuo L, et al. Myc increases self-renewal in neural progenitor cells through Miz-1. Journal of Cell Science. 2008;121(23):3941–3950.

48. Fults D, et al. MYC Expression Promotes the Proliferation of Neural Progenitor Cells in Culture and *In Vivo*. Neoplasia. 2002;4(1):32–39.

49. Zheng X, et al. Metabolic reprogramming during neuronal differentiation from aerobic glycolysis to neuronal oxidative phosphorylation. eLife. 2016;5:e13374.

50. Chau KF, et al. Downregulation of ribosome biogenesis during early forebrain development. eLife. 2018;7:e36998.

51. Wang J, et al. c-Myc Is Required for Maintenance of Glioma Cancer Stem Cells. PLOS ONE. 2008;3(11):e3769.

52. Zheng H, et al. p53 and Pten control neural and glioma stem/progenitor cell renewal and differentiation. Nature. 2008;455(7216):1129–1133.

53. Odia Y, et al. cMYC expression in infiltrating gliomas: associations with IDH1 mutations, clinicopathologic features and outcome. J Neurooncol. 2013;115(2):249–259.

54. Bai H, et al. Integrated genomic characterization of IDH1-mutant glioma malignant progression. Nat Genet. 2016;48(1):59–66.

55. Jenkins RB, et al. A low-frequency variant at 8q24.21 is strongly associated with risk of oligodendroglial tumors and astrocytomas with IDH1 or IDH2 mutation. Nat Genet. 2012;44(10):1122–1125.

56. Yanchus C, et al. A noncoding single-nucleotide polymorphism at 8q24 drives IDH1-mutant glioma formation. Science. 2022;378(6615):68–78.

57. Fu Y, et al. FTO-mediated formation of N6-hydroxymethyladenosine and N6-formyladenosine in mammalian RNA. Nat Commun. 2013;4(1):1798.

58. Murphey K, et al. Acute myeloid leukemia and dilated cardiomyopathy in a pediatric patient with D-2-hydroxyglutaric aciduria type I. American Journal of Medical Genetics Part A. 2022;188(9):2707–2711.

59. Srinivasan A, et al. IDH1 mutated acute myeloid leukemia in a child with metaphyseal chondromatosis with D-2-hydroxyglutaric aciduria. Pediatric Hematology and Oncology. 2020;37(5):431–437.

60. Yeetong P, et al. Widespread and debilitating hemangiomas in a patient with enchondromatosis and D-2-hydroxyglutaric aciduria. Skeletal Radiol. 2018;47(11):1577– 1582.

61. Hutton SR, Pevny LH. Isolation, Culture, and Differentiation of Progenitor Cells from the Central Nervous System. Cold Spring Harb Protoc. 2008;2008(11):pdb.prot5077.

62. Tasdogan A, et al. Metabolic heterogeneity confers differences in melanoma metastatic potential. Nature. 2020;577(7788):115–120.

63. DeVilbiss AW, et al. Metabolomic profiling of rare cell populations isolated by flow cytometry from tissues. eLife. 2021;10:e61980.

64. Aurora AB, et al. Loss of glucose 6-phosphate dehydrogenase function increases oxidative stress and glutaminolysis in metastasizing melanoma cells. PNAS. 2022;119(6). 10.1073/pnas.2120617119.

65. Su X, Lu W, Rabinowitz JD. Metabolite Spectral Accuracy on Orbitraps. Anal Chem. 2017;89(11):5940–5948.

66. Dobin A, Gingeras TR. Mapping RNA-seq Reads with STAR. Current Protocols in Bioinformatics. 2015;51(1):11.14.1–11.14.19.

67. Putri GH, et al. Analysing high-throughput sequencing data in Python with HTSeq 2.0. Bioinformatics. 2022;38(10):2943–2945.

68. Love MI, et al. RNA-Seq workflow: gene-level exploratory analysis and differential expression. F1000Res. 2016;4:1070.

69. Liu X, et al. *In Situ* Capture of Chromatin Interactions by Biotinylated dCas9. Cell. 2017;170(5):1028–1043.e19.

70. Langmead B, et al. Ultrafast and memory-efficient alignment of short DNA sequences to the human genome. Genome Biology. 2009;10(3):R25.

71. Zhang Y, et al. Model-based Analysis of ChIP-Seq (MACS). Genome Biology. 2008;9(9):R137.

