## Supplemental figures for "L-2-hydroxyglutarate impairs neuronal differentiation through epigenetic activation of *MYC* expression"

Supplemental Figure 1

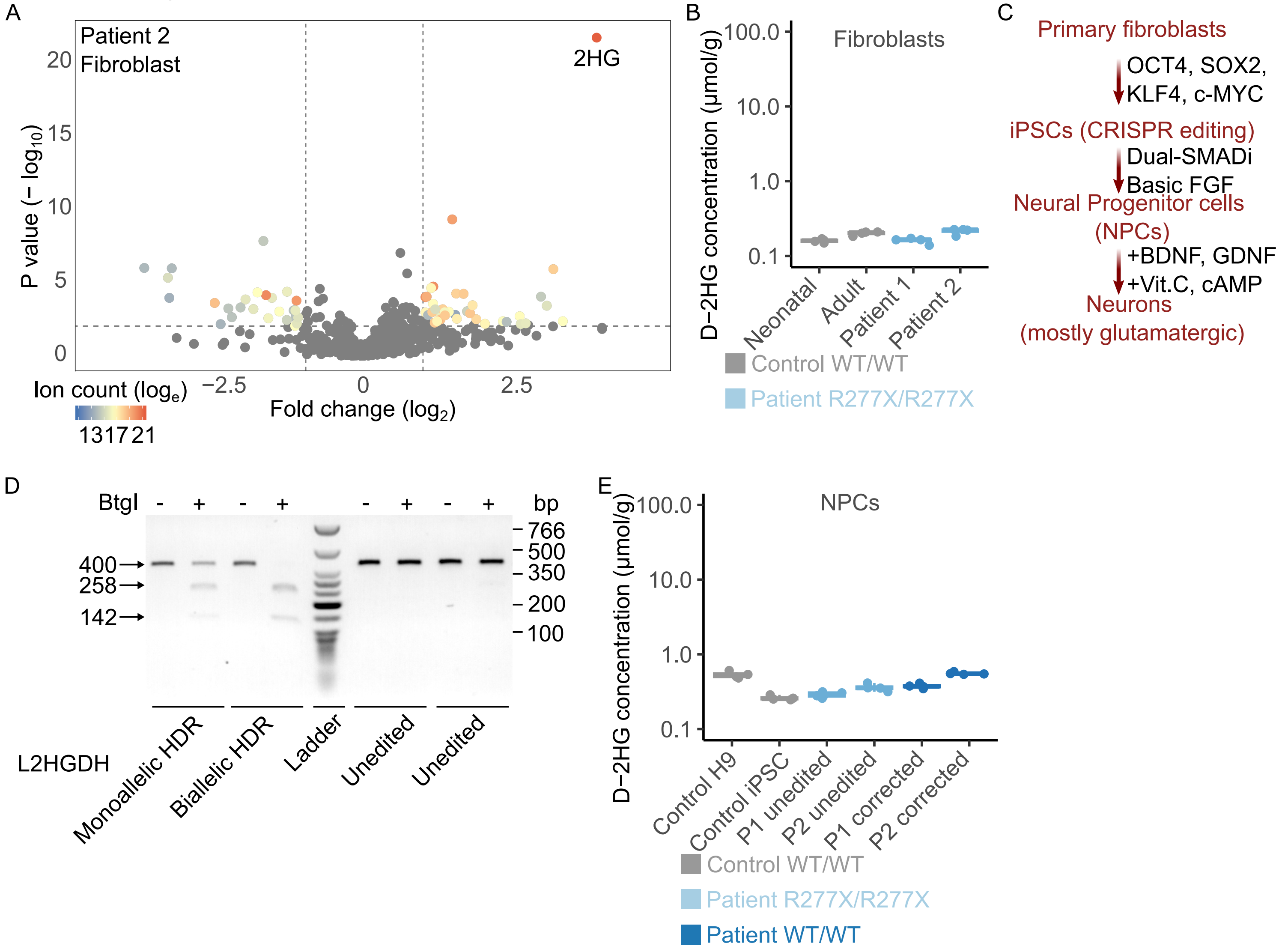

**Supplemental Figure 1. An isogenic, patient-derived iPSC system to study L2HGDH deficiency.**

(A) Volcano plot of fibroblast metabolites comparing Patient 2 to 29 unrelated lines (sibling excluded), each profiled in quadruplicate. A mixed-effects model was used as in Figure 1C, with cell line as the random effect.

(B) D-2HG concentrations in neonatal control fibroblasts, adult control fibroblasts, and L2HGA patient fibroblasts.

(C) Schematic of neuronal differentiation using patient-derived induced pluripotent stem cells (iPSCs), starting from patient fibroblasts and progressing through NPCs to mature neurons (predominantly glutamatergic).

(D) Restriction fragment length polymorphism (RFLP) analysis of iPSC clones following CRISPR/Cas9 genome editing, showing the digestion pattern with *BtgI* to differentiate between unedited, monoallelic HDR, and biallelic HDR clones.

(E) D-2HG concentrations in neural progenitor cells (NPCs) derived from various hPSC lines, including control H9 NPCs, control iPSC-derived NPCs, Patient 1 unedited NPCs, Patient 2 unedited NPCs, Patient 1 corrected NPCs, and Patient 2 corrected NPCs.

For (B) and (E), data are shown as box plots with jittered individual data points. Boxes indicate the interquartile range (25th–75th percentile), with the horizontal line marking the median. Whiskers extend to the minimum and maximum values within 1.5× the interquartile range. Statistical significance was assessed using one-way ANOVA followed by Tukey's HSD test ($n$ = 4 per group).

Supplemental Figure 2

A

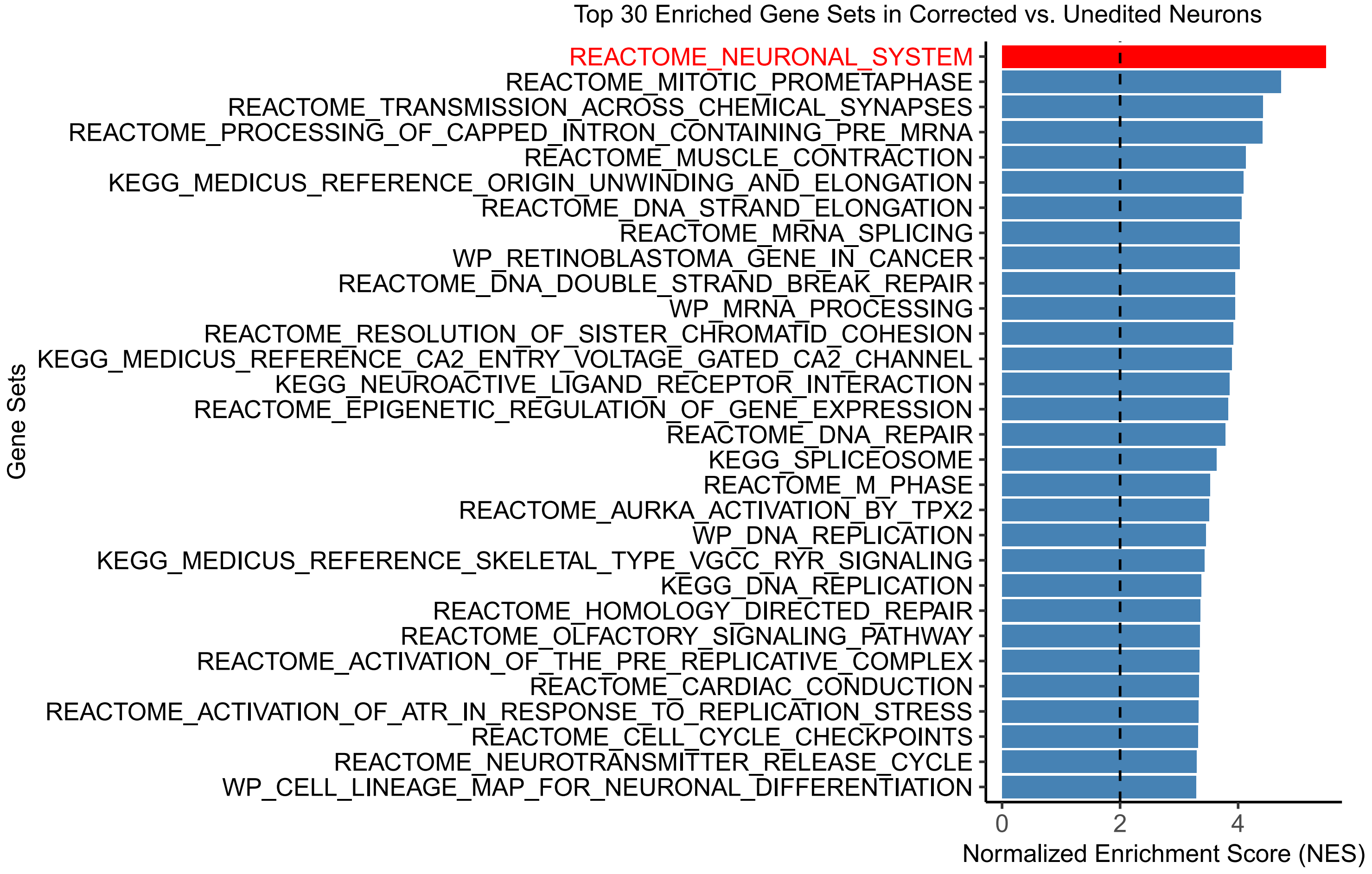

**Supplemental Figure 2. L-2HG accumulation enhances NPC proliferation and self-renewal while inhibiting neuronal differentiation.**

(A) Top 30 enriched gene sets from gene set enrichment analysis (GSEA) comparing corrected neurons vs. unedited neurons. The X-axis represents the normalized enrichment score (NES), and rows indicate the top 30 enriched gene sets. The dashed line at NES = 2.0 marks a commonly used threshold for significant enrichment. All 3917 Canonical Pathways (c2.cp) gene sets from the Human MSigDB Collections were used for GSEA.

Supplemental Figure 3

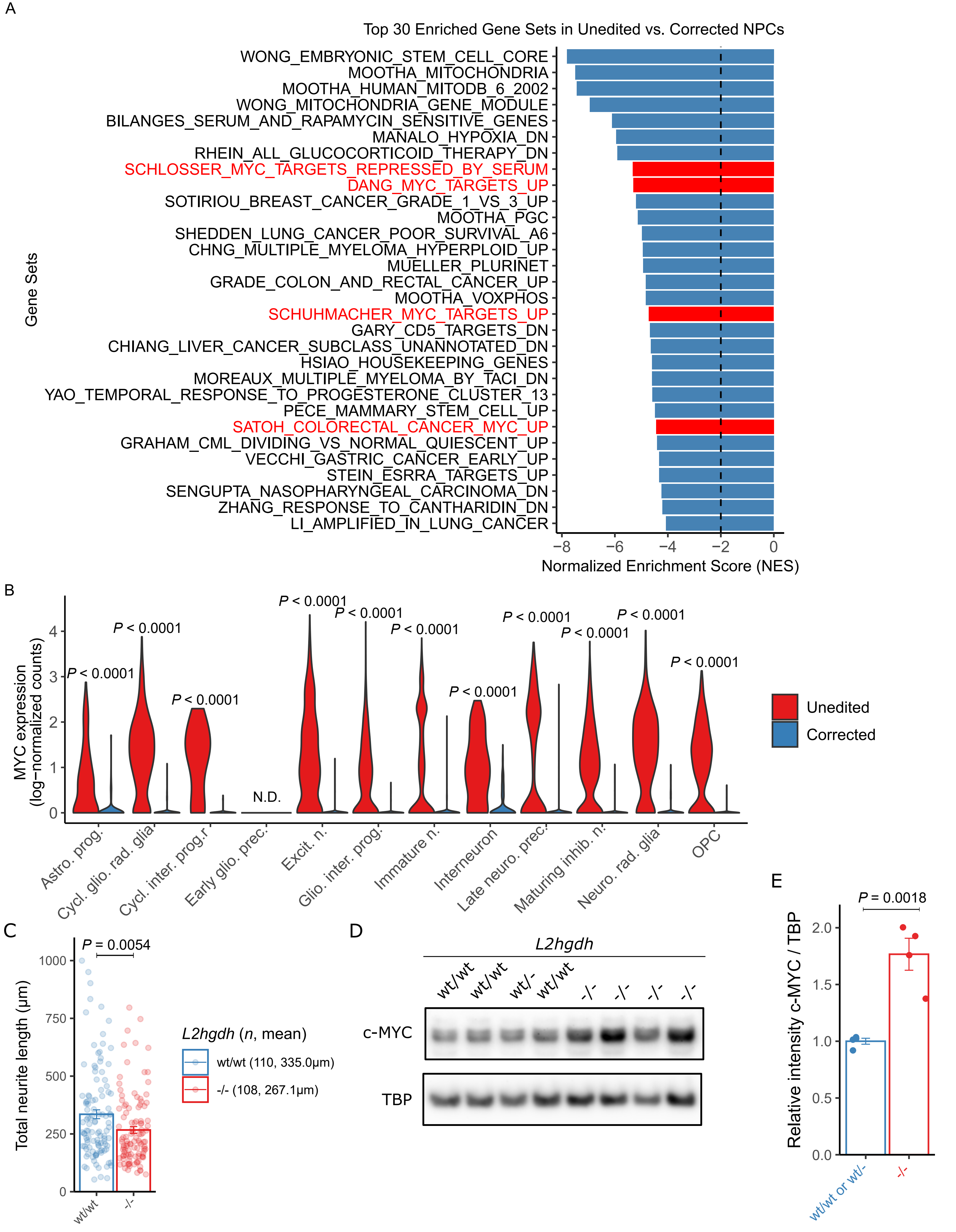

**Supplemental Figure 3. Increased *MYC* Expression in L2HGDH-Deficient NPCs.**

(A) Top 30 enriched gene sets from gene set enrichment analysis (GSEA) comparing corrected vs. unedited NPCs. The X-axis represents the normalized enrichment score (NES), and rows indicate the top 30 enriched gene sets. The dashed line at NES = 2.0 marks a commonly used threshold for significant enrichment. All 3,494 Chemical and Genetic Perturbations (c2.cgp) gene sets from the Human MSigDB Collections were used for GSEA.

(B) Violin plots showing *MYC* expression (log-normalized counts) in unedited and corrected cells from day 45 human cortical spheroids described in Figure 2K.

(C) Quantification of total neurite lengths in wild-type (*wt/wt*) and *L2hgdh-/-* neurons derived from mouse NPCs.

(D) Immunoblot analysis of nuclear c-Myc in wild-type (*wt/wt*) and *L2hgdh-/-* NPCs derived from E15.5 mouse embryos. TBP was used as a loading control for nuclear lysates.

(E) Quantification of relative c-Myc levels, normalized to TBP, as described in (D).

Error bars represent ± 1 SEM of biological replicates. For (B), expression values reflect normalized, log-transformed transcript counts. For each cell type, statistical significance was assessed using a two-sided Wilcoxon rank-sum test. *P* values were adjusted for multiple comparisons using the Benjamini-Hochberg method. Statistical significance was determined using Student’s t test (C and E). For (C), sample sizes and mean neurite lengths are indicated in the figure; for (D), $n = 4$.

Supplemental Figure 4

A

| Histone Mark | FDR | Ranking |
|---|---|---|
| H3K4me2 | 1.33E-22 | 932 out of 11,259 |
| H3K4me3 | 3.00E-06 | 335 out of 5,977 |

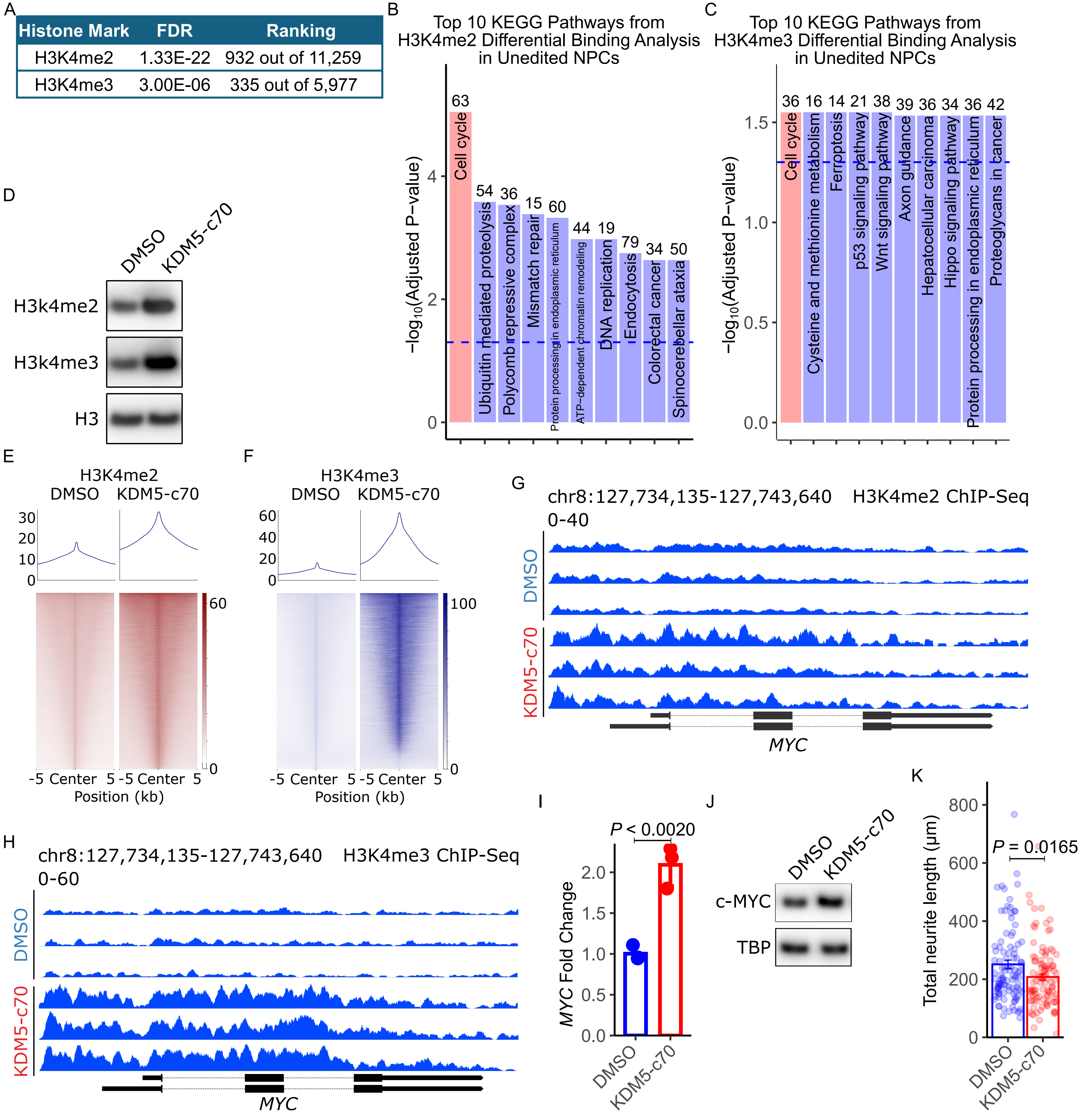

**Supplemental Figure 4. Increased activating histone methylation marks H3K4me2 and H3K4me3 at the *MYC* locus in L2HGDH-Deficient NPCs.**

(A) Statistical significance of differential binding analysis for H3K4me2 and H3K4me3 markers, comparing unedited vs. corrected Patient 1 NPCs. The adjusted false discovery rate (FDR) ranking of each histone mark is shown. The third column shows the ranking of the MYC locus among all differential loci.

(B) and (C) Top 10 KEGG pathways identified from differential binding analysis of H3K4me2 (B) and H3K4me3 (C) in unedited vs. corrected Patient 1 NPCs, based on ChIP-seq data. The Y-axis represents the $-\log_{10}$ (*P*-value) of pathway enrichment, while the columns indicate the top 10 enriched pathways. Numbers above the bars indicate the number of genes associated with each pathway. The blue dashed line marks the significance threshold at $P = 0.05$.

(D) Immunoblot analysis of activating histone marks H3K4me2 and H3K4me3 in corrected Patient 1 NPCs treated with DMSO or the KDM5 inhibitor C70 (25 μM) for 14 days. Histone H3 was used as a loading control for histone extraction.

(E) and (F) ChIP–Seq profiles of H3K4me2 (E) and H3K4me3 (F) in corrected Patient 1 NPCs treated with DMSO or the KDM5 inhibitor C70 (25 μM) for 14 days. ChIP–Seq signals were plotted over center peaks (± 5 kb from peak center) identified in DMSO-treated NPCs. Sites were sorted by the ChIP–Seq signal intensity from DMSO-treated NPCs.

(G) and (H) Representative ChIP-seq tracks showing H3K4me2 (G) and H3K4me3 (H) enrichment at the *MYC* locus in corrected Patient 1 NPCs treated with DMSO or the KDM5 inhibitor C70 (25 μM) for 14 days.

(I) qRT-PCR analysis of *MYC* mRNA levels (normalized to 36B4) in corrected Patient 1 NPCs treated with DMSO or KDM5 inhibitor C70 (25 μM) for 14 days. Statistical analysis was performed to compare qPCR fold change values between the DMSO and C70 groups. First, data normality was assessed using the Shapiro-Wilk test, confirming that both groups followed a normal distribution ($P > 0.05$). Based on this, an independent t-test was applied, yielding a statistically significant result ($P$ = 0.0020). $n$ = 3 biological replicates.

(J) Immunoblot analysis of nuclear c-MYC levels in corrected Patient 1 NPCs treated with DMSO or the KDM5 inhibitor C70 (25 μM) for 14 days. TBP was used as a loading control for nuclear lysates.

(K) Quantification of total neurite lengths in corrected Patient 1 NPCs treated as described in (J). Neurite lengths were measured using the SNT plugin in ImageJ from immunofluorescent images of fixed cells. Data are presented as box plots with jittered individual data points. Boxes represent the interquartile range (25th–75th percentile), the horizontal line indicates the median, and whiskers extend to values within 1.5× the interquartile range. Statistical significance was determined by Student's t-test (n = 105 per group).

Supplemental Figure 5

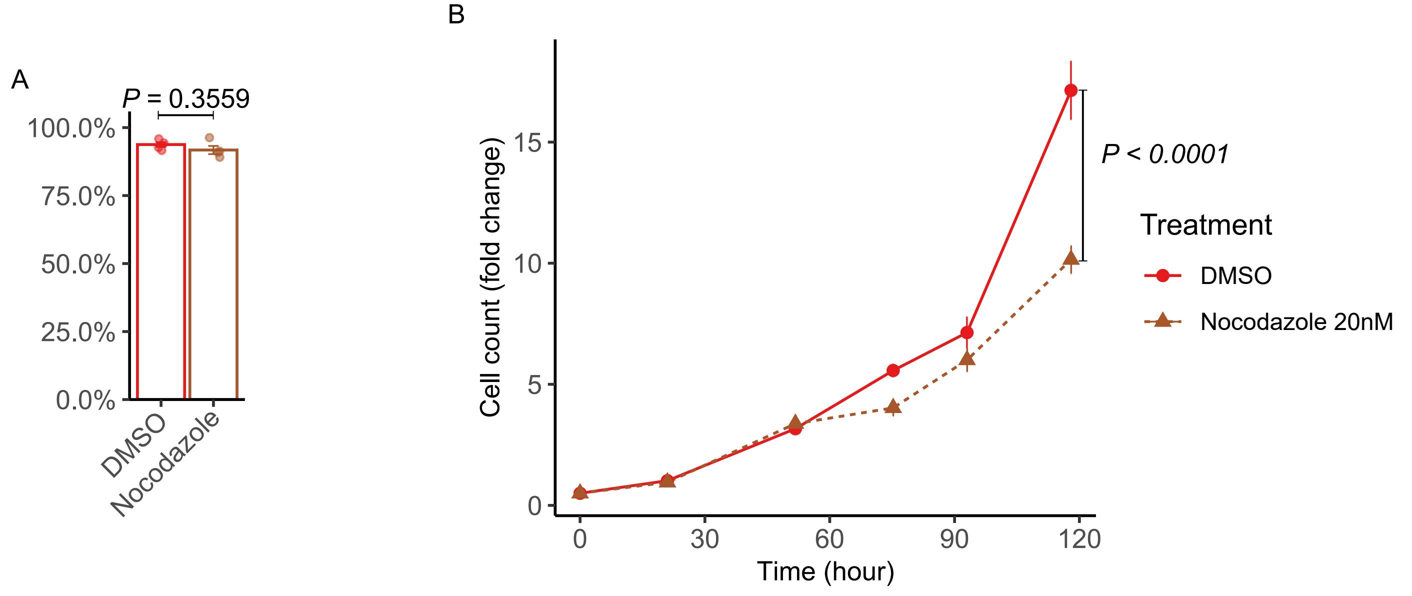

**Supplemental Figure 5. Nocodazole treatment of unedited NPCs.**

(A) Cell viability analysis of unedited Patient 1 NPCs treated with DMSO or 20 μM nocodazole for 7 days. Cell viability was measured as a percentage of total viable cells in each condition. Error bars represent ± 1 SEM of three biological replicates. Statistical significance was determined using Student's t test.

(B) Growth curves of unedited Patient 1 NPCs treated with DMSO or 20 μM nocodazole. Error bars represent ±1 SEM of biological replicates. Differences in cell growth over time were assessed using a two-way ANOVA on fold-change data, with treatment and time point as independent factors. Three biological replicates were included for each condition at each time point, with an assumption of independent sampling across groups
